# Kinship estimation bias carries over to heritability estimation bias using variance components

**DOI:** 10.1101/2025.05.13.653659

**Authors:** Zhuoran Hou, Alejandro Ochoa

## Abstract

Heritability is a fundamental parameter of diseases and other traits, quantifying the contribution of genetics to that trait. Kinship matrices, also known as Genetic Relatedness Matrices or “GRMs”, are required for heritability estimation with variance components models. However, the most common “standard” kinship estimator employed by GCTA and other approaches, can be severely biased in structured populations. In this study, we characterize heritability estimation biases in GCTA due to kinship estimation biases under population structure. For the standard *ratio-of-means* (ROM) kinship estimator, we derive a closed-form expression for heritability bias given by the mean kinship value and the true heritability. The standard *mean-of-ratios* (MOR) estimator is the most widely used in practice, and exhibits more severe bias than ROM due to upweighing low-frequency variants. Using simulation studies with admixture and family structures, as well as simulated traits from 1000 Genomes genotypes, we find that only Popkin, which is the only unbiased population kinship estimator, produces unbiased heritability estimates in structured settings. Pedigree-only estimates have upward heritability biases when there is population structure. Finally, we analyze three structured datasets with real phenotypes—the San Antonio Family Study, the Hispanic Community Health Study / Study of Latinos, and a multiethnic Nephrotic Syndrome cohort. The standard MOR estimator can produce both downward and upward heritability biases depending on population structure and variant frequency spectrum, compared to the other two estimators. Overall, common kinship estimators result in heritability estimation biases when applied to structured populations, a challenge that Popkin successfully overcomes.

## 1 Introduction

Heritability is an important parameter of diseases and other traits, quantifying the contribution of genetics to that trait as opposed to non-genetic environmental factors (Lush et al., 1949). Heritability is reflected in the extent to which relatives have similar phenotypes, which is the reason relatedness plays a large role in heritability estimation (Visscher et al., 2008). In addition, heritability is an important parameter of many trait models, including polygenic risk scores (PRS), whose performance is bounded above by heritability (Choi et al., 2020). There are two primary types of heritability: broad-sense heritability (*H*^2^) and narrow-sense heritability (*h*^2^). Broad-sense heritability includes all genetic variance components—additive (A), dominance (D), and epistatic (I) effects—while narrow-sense heritability focuses specifically on additive genetic variance (Falconer, 1996). Because additive effects are transmitted from parent to offspring in a predictable manner, narrow-sense heritability is the relevant parameter most often estimated in GWAS, PRS modeling, and many other applications. Accurately estimating *h*^2^ is therefore fundamental for both understanding genetic contributions to traits and for the development of predictive models.

Heritability has long been estimated from close relatives, such as twins or siblings (Falconer, 1996). The variance component model of SOLAR, which is based on estimating kinship matrices from pedigrees, enables the use of more complex pedigrees and distant relatives (Almasy and Blangero, 1998a). GCTA extended the last approach to population data, employing population kinship matrices estimated from genetic data only, which was used to demonstrate that SNPs likely explain the majority of missing heritability for height (Yang et al., 2010; Yang et al., 2011). Lastly, there are alternate approaches for estimating heritability from GWAS summary statistics and LD estimates, which enable partitioning heritability within gene sets, such as LD Score Regression (Bulik-Sullivan et al., 2015; Luo et al., 2021) and SumHer (Speed and Balding, 2019), but since they do not use kinship matrices they fall out of the scope of the present work.

Despite widespread use, all heritability estimation methods face important limitations. Traditional twin studies, for example, assume that monozygotic and dizygotic twins share environments to the same extent—a questionable assumption that can inflate heritability estimates by conflating genetic and environmental similarity (Tenesa and Haley, 2013; Charney, 2012). Pedigree-based methods like SOLAR are sensitive to incomplete or biased family structures. GCTA, although groundbreaking, has also sparked debate: it primarily captures the additive effects of common SNPs, potentially missing contributions from rare variants, non-additive effects, or poorly tagged genomic regions (Speed et al., 2012a; Speed et al., 2017a; Zaitlen et al., 2013; Yang et al., 2017). Moreover, population structure and cryptic relatedness can bias GCTA-based estimates if not properly controlled, especially in admixed samples (Price et al., 2010; Yang et al., 2017; Huang et al., 2025). These issues have led to an ongoing reevaluation of how heritability is conceptualized and measured in the genomic era.

Accurate kinship estimation is crucial for heritability estimation based on variance components, such as GCTA, which are special cases of linear mixed-effects models (LMMs). However, the most common kinship estimator employed by these approaches, which we refer to as the “standard” estimator, can be severely biased in structured populations (Ochoa and Storey, 2021). We previously showed that association tests are invariant to the use of common biased kinship estimators compared to an unbiased estimator (Hou and Ochoa, 2023). However, heritability estimation requires unbiased estimates of the random effect coefficient, which is biased when the standard kinship estimator is used. Thus, we hypothesize kinship estimation bias will have a strong effect on heritability estimation accuracy, particularly when there is population structure. However, we do not expect these biases to explain missing heritability in previous studies of relatively homogeneous populations, such as the landmark height paper which only analyzed Australian individuals of European ancestry (Yang et al., 2010).

In addition to concerns about population structure, recent efforts have sought to improve heritability estimation by modeling the effects of rare variants through partitioned genomic relationship matrices (GRMs) or by adjusting assumptions about genetic architecture. Several studies have emphasized that the contribution of a variant to heritability depends not only on its effect size but also on its minor allele frequency (MAF) and linkage disequilibrium (LD) patterns. Our theory suggests that these differences could be in part due to kinship estimation biases specific to rare variants. However, another potentially complementary explanation is that rare variants have a different structure than common variants in humans (Zaidi and Mathieson, 2020). Traditional methods overestimate the contribution of common compared to rare variants, and it has been proposed that heritability is more evenly distributed across the MAF spectrum when accounting for LD (Speed et al., 2017a). Furthermore, low-frequency variants are enriched for functional annotations under negative selection, reinforcing the view that rare variants often have larger per-allele effects (Gazal et al., 2018). Using whole-genome sequencing data, it has been shown that rare variants contribute substantially to the heritability of complex traits—possibly explaining a significant portion of the missing heritability (Wainschtein et al., 2022). More flexible models have been introduced that accommodate diverse genetic architectures, including skewed effect size distributions correlated with allele frequency, but still found that rare variant heritability remains difficult to estimate accurately (Hou et al., 2019). SumHer models effect sizes as a function of both LD and MAF, producing more realistic assumptions about rare variants compared to GCTA (Speed and Balding, 2019). Taken together, these studies highlight the importance of modeling rare variant contributions explicitly, both in terms of effect-size distributions and their relationships to allele frequency and LD, which are essential for accurate heritability estimation in complex trait genetics. However, our work highlights important unsolved challenges estimating variance for rare variants, particularly under population structure, which remain an open problem.

In this study, we characterize the theoretically predicted heritability estimation bias due to kinship bias, following the derivation in our previous work (Hou and Ochoa, 2023) and others (Chen and Storey, 2022), and empirically compare key kinship estimators in various population structure scenarios. We conduct simulation studies to evaluate heritability estimation accuracy and bias under scenarios such as admixture structure only and admixture combined with family structure, as well as simulated traits drawn from the real 1000 Genomes genotypes. We then apply these kinship estimators to datasets with population structure and real phenotypes, to further characterize the sources and extent of bias in practice: the San Antonio Family Study (Mitchell et al., 1996), the Hispanic Community Health Study/Study of Latinos (Sorlie et al., 2010), and a Nephrotic Syndrome multiethnic cohort. Our results show that the standard kinship estimators—particularly the commonly used mean-of-ratios (MOR) version—introduce systematic and sometimes severe biases in heritability estimation, especially in the presence of population structure and rare variants. On the other hand, pedigree-derived kinship matrices exhibit upward biases when there is population structure. Only the Popkin estimator yields unbiased heritability estimates across all simulation and real data settings. Overall, our findings highlight the importance of using unbiased kinship estimators to obtain reliable heritability estimates in structured populations.

## 2 Methods

### 2.1 Genetic and trait models

Suppose that there are *m* biallelic loci and *n* diploid individuals. The genotype *x*_*ij*_ ∈ {0, 1, 2} at a locus *i* of the individual *j* is encoded as the number of reference alleles, for a pre-selected but otherwise arbitrary reference allele per locus. Let *φ*_*jk*_ is the kinship coefficient of two individuals *j* and *k*, and *p*_*i*_ is the ancestral allele frequency at locus *i*, which are in terms of an implicit ancestral population (usually the most recent common ancestor) that will remain fixed in this work. Under the kinship model (Malécot, 1948; Wright, 1949; Jacquard, 1970; Astle and Balding, 2009; Ochoa and Storey, 2021) the expectation and covariance of genotypes are given by

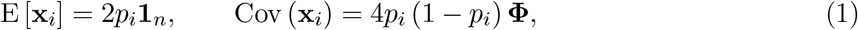

where **x**_*i*_ = (*x*_*ij*_) is the length-*n* vector of genotypes at locus *i*, **Φ** = (*φ*_*jk*_) is the *n*×*n* kinship matrix, and **1**_*n*_ is a length-*n* vector of ones. The definition of kinship that we follow, which is a probability of identity by descent, is such that the maximum kinship of 1 is achieved for fully inbred individuals (who have only homozygote genotypes) in the case of self-kinship, or fully inbred identical twins for a pair of different individuals. In this definition, the self kinship of an outbred individual is 1/2, the kinship between a parent and their outbred child is 1/4, and so is the expected kinship between outbred siblings. In contrast, GCTA and related models typically define kinship as twice the value that we use, so that self kinship for an outbred individual is 1, and kinship between outbred parent and child or between siblings is 1/2 (Yang et al., 2010; Yang et al., 2011).

The quantitative trait vector **y** for all individuals is assumed to follow a linear polygenic model,

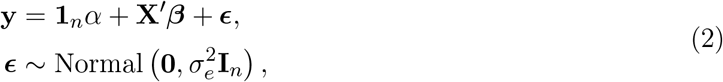

where *α* is the intercept coefficient, ***β*** = (*β*_*i*_) is a length-*m* vector of genetic effect coefficients for each locus *i*, ***ϵ*** is a length-*n* vector of non-genetic independent residual effects with zero mean and standard deviation 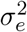, which is also called the environmental variance component, **I**_*n*_ is the *n* × *n* identity matrix, and the prime (^′^) denotes matrix transposition.

We now derive the variance component model of interest from the previous genotype and trait models. As stated in those models, **X** and ***ϵ*** are random, while we treat *α* and ***β*** as fixed parameters. Note that the mean trait is given by

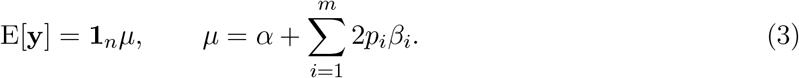

Denote the genetic effect by **s** = **X**^′^***β***. The covariance structure of the genetic effect is also a scaled version of the kinship matrix. In particular, assuming Eq. (1) and independent loci, then

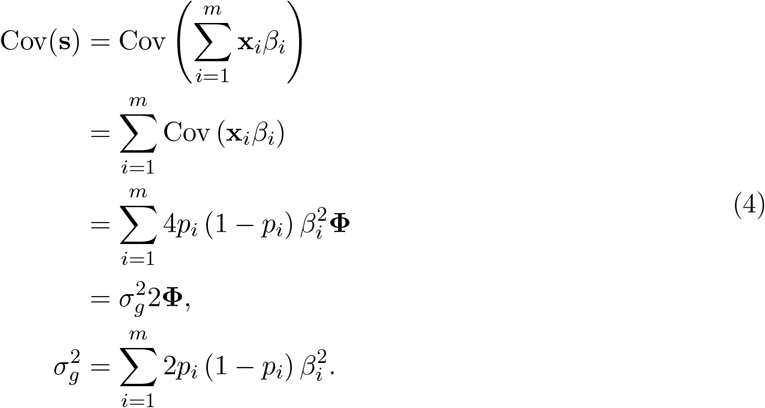

This particular scale for the genetic variance component 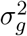, which leaves a factor of two behind, is used traditionally so that 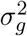 corresponds to the variance of outbred individuals, who have *φ*_*jj*_ = 1*/*2. Further, as noted earlier, GCTA and other models define their kinship matrices as 2**Φ** under our notation.

Implicit in the assumed Eq. (1) and the resulting Eq. (4) is that the covariance structure is the same for causal variants as for the rest of the genome (from which the kinship matrix will be estimated, as shown later). However, there is evidence for humans that rare variants have a different structure than common variants (Zaidi and Mathieson, 2020), a potential misspecification to keep in mind as we analyze the real data later.

If we shift the mean of genotypes to the intercept, and assume that **s** is well approximated by a multivariate distribution with the above covariance, then we arrive at the model that GCTA fits:

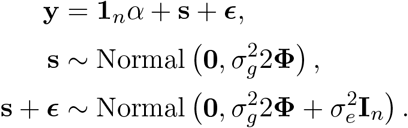

The narrow-sense heritability *h*^2^ is defined as the proportion of variance that corresponds to the genetic variance component:

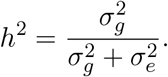

If we define the trait variance scale as 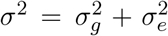, which is the trait variance of an outbred individual, then note that 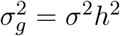 and 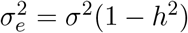.

### 2.2 Statistical problems of rare variant allele frequency estimates

A recurrent theme in this work, surfacing in both the biases of kinship estimators and of trait simulations, is statistical problems due to estimation of allele frequencies under population structure. The standard ancestral allele frequency estimator,

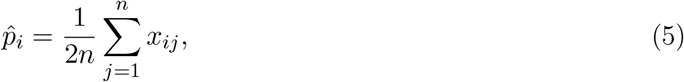

is unbiased 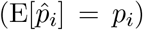 under the kinship model of Eq. (1), and has a variance of (Ochoa and Storey, 2021)

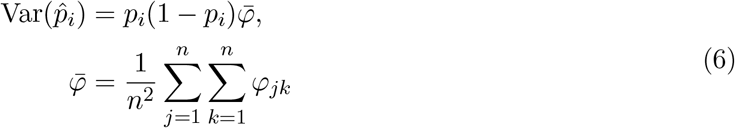

where 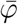 is the mean kinship value in the sample, and converges to a non-zero value when there is population structure. For this reason, 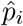 is not a consistent estimator when there is population structure, in other words it does not converge to the true *p*_*i*_ as sample sizes go to infinity, which is an underlying assumption of many previous works in statistical genetics that does not hold for real data with population structure.

This work requires estimates of the Binomial variance factor *p*_*i*_(1 − *p*_*i*_). Previous work found that the sample estimator is biased, although it can be readily unbiased with a good estimate of 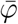 (Ochoa and Storey, 2021):

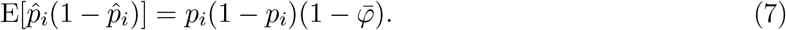

However, using a normality approximation, in Appendix A we prove that the variance of this estimator also converges to a non-zero value, so 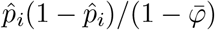 is also not consistent:

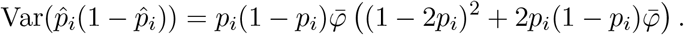

As a direct consequence, the standard estimate of the inverse Binomial variance can be severely biased. Specifically, applying Jensen’s inequality to the inverse function with positive arguments, combined with the unbiased form that follows from Eq. (7), we obtain that

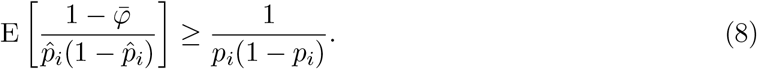

Equality occurs when 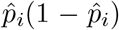 has no variance, which again does not occur under population structure, and conversely, higher variance exacerbates the inequality. Empirically, biases are greater for rare variants, where *p*_*i*_ or 1 − *p*_*i*_ are close to zero.

### 2.3 Kinship estimation

Each estimator bias type has two locus weight types called *ratio-of-means* (ROM) and *mean-of-ratios* (MOR) (Bhatia et al., 2013; Ochoa and Storey, 2021). Only ROM estimators have closed-form limits. Let 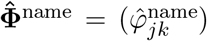 relate the scalar and matrix formulas of each named kinship estimator.

#### 2.3.1 Standard kinship estimator

The standard kinship estimator is the most widely used estimator in various applications of population structure (Astle and Balding, 2009; Speed and Balding, 2015; Wang et al., 2017a), including heritability estimation (Yang et al., 2010; Yang et al., 2011; Speed et al., 2012b; Speed and Balding, 2015; Speed et al., 2017b) and genetic association tests based on PCA (Price et al., 2006), LMM (Astle and Balding, 2009; Zhou and Stephens, 2012; Loh et al., 2015; Sul et al., 2018), and other models (Rakovski and Stram, 2009; Thornton and McPeek, 2010).

The ROM and MOR versions of the standard kinship estimator are, respectively,

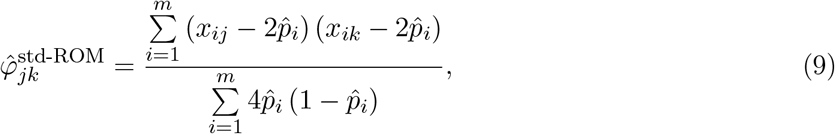

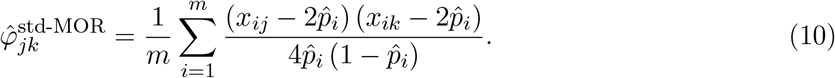

The ROM estimator has a biased limit (Ochoa and Storey, 2021; Hou and Ochoa, 2023):

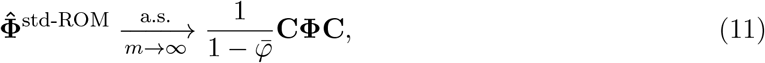

where **Φ** is the true kinship matrix, the scalar mean kinship 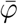 is as in Eq. (6), and 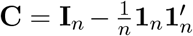 is the centering matrix. The MOR estimator does not have a closed-form limit, but in practice it is well approximated by Eq. (11) when rare variants are excluded prior to calculating this estimate, though they can differ greatly otherwise due to the additional biases described in Eq. (8).

#### 2.3.2 Popkin kinship estimator

The popkin (population kinship) estimator (Ochoa and Storey, 2021) is given by

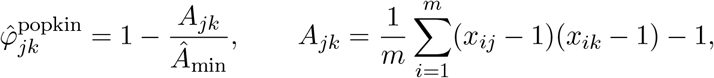

where 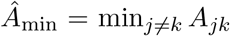. This estimator of type ROM has an unbiased almost sure limit as the number of loci *m* goes to infinity,

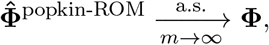

under the assumption that the true minimum kinship is zero. Popkin avoids the biases of the standard estimators because it does not rely on the inconsistent 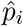 estimator.

The mean kinship values 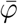 reported here are always calculated from the true kinship matrix, or estimated with popkin when the true kinship matrix is unavailable, as is the case for real datasets, since popkin estimates this value without bias. Note both Standard ROM and MOR mean kinship estimates are always exactly zero (algebraically, with no variance), which indirectly confirms that they must be biased.

#### 2.3.3 Software

Standard MOR kinship matrices and all heritability estimates are calculated using GCTA (version 1.93.2beta) (Yang et al., 2011). Popkin kinship estimates are computed using the `popkin` R package, while Standard ROM kinship estimates are calculated using the `popkinsuppl` R package (Ochoa and Storey, 2021). Kinship matrices are calculated from pedigrees using the `kinship2` R package (Sinnwell et al., 2014). GCTA is provided alternative kinship matrices encoded as standard GRM files using the `genio` R package (Yao and Ochoa, 2022), doubling kinship values to match Eq. (4) as needed. Plink (version 2.00a3LM) is used to process genotype data (Chang et al., 2015).

### 2.4 Heritability estimation bias due to Standard ROM kinship bias

We recently characterized the theoretical relationship between variance component estimates of a biased kinship matrix and its unbiased counterpart (Hou and Ochoa, 2023). In particular, when provided with the Standard ROM kinship matrix of Eq. (11), the LMM fits the genetic variance component with an algebraically biased form compared to the unbiased estimate from Popkin, whereas the environment variance component estimate is the same for both:

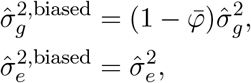

where 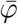 is the mean value of the true kinship matrix of Eq. (6). Therefore, since 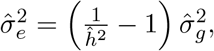, Standard ROM heritability estimates will be biased with the following form, where 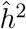 is the unbiased Popkin estimate:

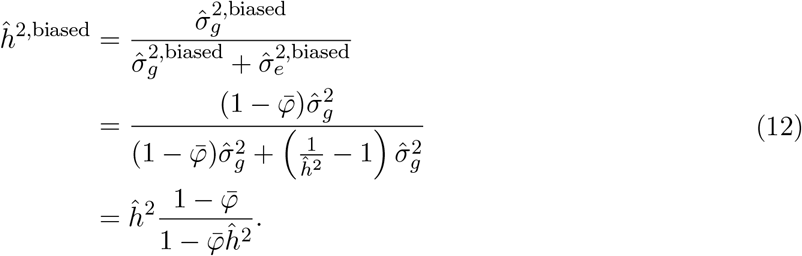

The above formula not only relates Popkin and Standard ROM estimates in a given dataset, but it also holds in their limit, in which case the Popkin estimate becomes the true heritability. In terms of estimate limits, bias is larger for intermediate true heritability estimates and increases with 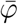, while bias decreasing to zero when the heritability is close to 0 or 1 (Fig. 1).

**Figure 1:**
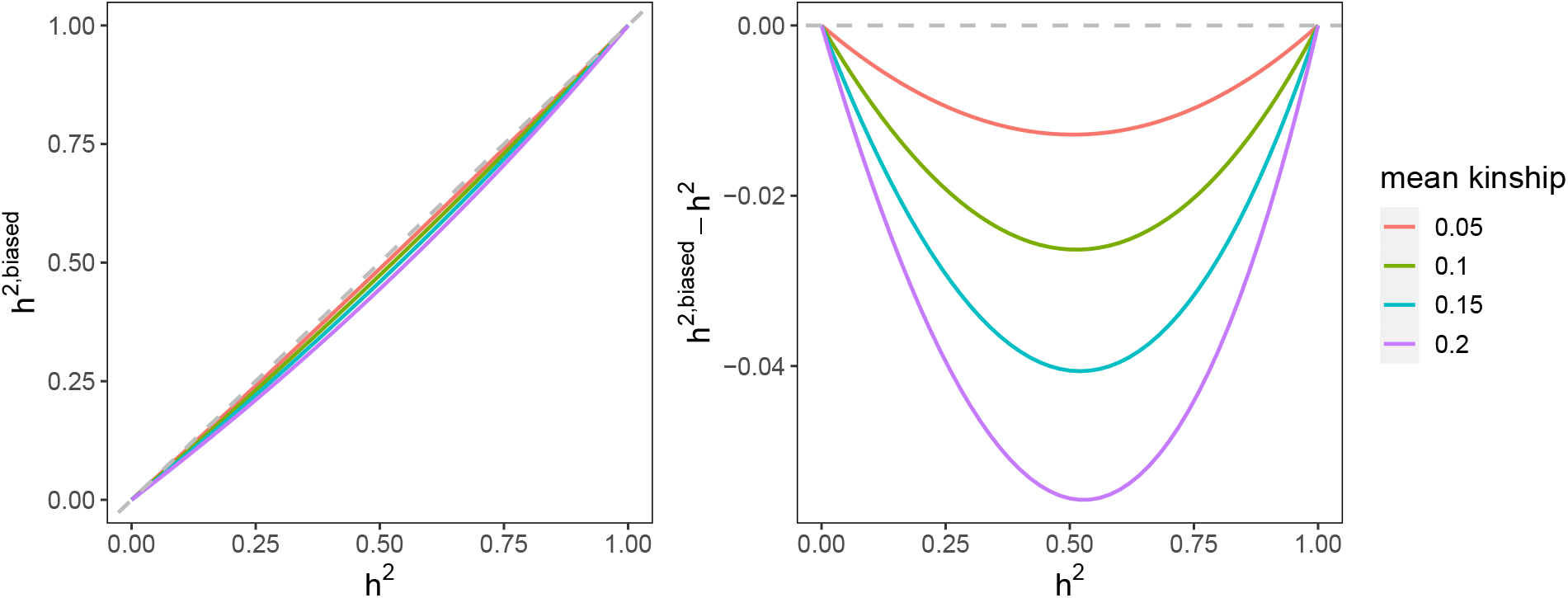
Relationship between the true heritability and biased Standard ROM estimates. We illustrate the functional form of Eq. (12) for a range values of the mean kinship parameter 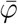 observed in real human datasets (Table 1). Note Standard MOR (the most commonly used estimator) has even larger biases empirically. Left: the relationship between biased estimates and the true heritability. Right: the relationship between hertability bias and true heritability.

### 2.5 Simulations

To characterize the effect of kinship estimator bias in heritability estimation, we use simulated genotypes and traits and estimate kinship matrices from these genotypes. Each scenario was replicated 50 times, in each case producing a new genotype matrix.

**Table 1:** Proportions of rare variants and mean kinship values in the simulated and real datasets.

| Dataset | QC Filter | Proportion of loci with $MAF \leq 0.05$ | Mean kinship |
| --- | --- | --- | --- |
| Admix Sim. | $MAC > 0$ | 0.194 | 0.150 |
| Admix+Fam Sim. | $MAC > 0$ | 0.180 | 0.163 |
| 1000 Genomes | $MAC > 0$ | 0.930 | 0.133 |
| 1000 Genomes | $MAF \geq 0.01$ | 0.438 | 0.134 |
| 1000 Genomes | $MAF \geq 0.05$ | 0.000 | 0.100 |
| HCHS/SOL | $MAF \geq 0.01$ | 0.256 | 0.108 |
| SAMAFS | $MAF \geq 0.01$ | 0.281 | 0.063 |
| NS | $MAC \geq 20$ | 0.564 | 0.124 |

#### 2.5.1 Admixture simulation for genotype matrices

An admixed family is simulated as before (Yao and Ochoa, 2022; Hou and Ochoa, 2023), with *K* = 3 ancestries and *F*_ST_ = 0.3 for the admixed individuals, which resembles Hispanics and African Americans. Briefly, our admixture model simulates *n* = 1000 individuals with *m* = 100, 000 loci. Random ancestral allele frequencies 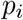 for *i* ∈ {1, …, *m*}, subpopulation allele frequencies 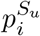 for *u* ∈ {1, …, *K*}, and individual-specific allele frequencies *π*_*ij*_ and genotypes *x*_*ij*_ for *j* ∈ {1, …, *n*} are drawn from this hierarchical model:

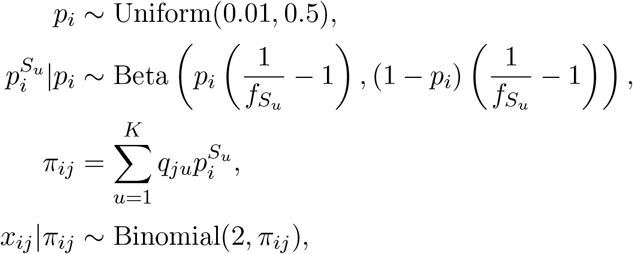

where this Beta is the Balding-Nichols distribution (Balding and Nichols, 1995) with mean *p*_*i*_ and variance 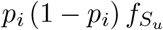. The admixture proportions 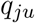 and subpopulation inbreeding values 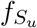 are constructed based on a diffusion model on a 1D geography (Ochoa and Storey, 2021).

For simulations with family structure, 20 generations are generated iteratively, as follows. Individuals in the first generation (*n* = 1000) are drawn from our admixture model, ordered by 1D geography, randomly assigned sex, and treated as locally unrelated. From subsequent generations, individuals are paired iteratively: randomly choosing males from the pool and pairing them with the nearest available female with local kinship *<* 1*/*4^3^ (to preserve the admixture structure) until there are no available males or females. Family sizes are drawn randomly ensuring every family has at least one child. Children are reordered by the average coordinates of their parents, their sex are assigned randomly, and their alleles are drawn from parents independently per locus. The simulation is implemented in the R package `simfam`.

#### 2.5.2 Trait simulation algorithms

We wish to simulate a quantitative trait **y** that follows the linear polygenic model of Eq. (2), assuming the genotype matrix **X** is given, but the intercept *α* and the genetic coefficients ***β*** have not been determined, and they can be random, but they must result in *m*_1_ number of causal loci (with non-zero coefficients), a trait mean of *µ*, variance scale of *σ*^2^, and a heritability of *h*^2^. In this study, we set *m*_1_ = 500, *µ* = 0, *σ*^2^ = 1, *h*^2^ ∈ {0, 0.1, …, 1} for the admixture simulations and *h*^2^ = 0.8 for 1000 Genomes. The algorithms require either true ancestral allele frequencies *p*_*i*_ (available for simulations but never for real genotype data) or both its sample estimate 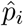 from Eq. (5) and an unbiased estimate of the mean kinship 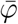 in order to unbias estimates using Eq. (7). These algorithms have been used to evaluate GWAS methods (Yao and Ochoa, 2022; Hou and Ochoa, 2023), but here we motivate them for evaluating heritability estimation.

In all cases, independent residual effects ***ϵ*** are simulated from Eq. (2) with 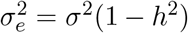, we randomly select *m*_1_ loci to be causal, and reindex loci to only include causal loci. Afterwards, the following paragraphs describe how to construct ***β*** under two evolutionary models, followed by the construction of *α*. In turn, each of those models has two cases, namely whether the true ancestral allele frequencies *p*_*i*_ are known or not.

In this paper, we treat effect sizes *β*_*i*_ as random variables only in the context of trait simulation (RC and FES), where they are explicitly sampled from a distribution to reflect the assumed genetic architecture. This randomness is necessary for defining and analyzing the properties of our estimators. However, in the rest of the paper we treat the *β*_*i*_ as fixed parameters. This separation maintains both the statistical rigor of our simulation-based proofs and the interpretability expected in applied genetics. In the simulation procedure, we explicitly track both the random variables defined by the model and their realized values in each replicate.

##### Random Coefficients (RC) model

The initial effect sizes *β*_*i*0_ are treated as independent random variables drawn from a standard normal distribution:

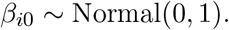

Let *b*_*i*0_ denote the realized value of *β*_*i*0_ used in the simulation procedure.

Then, if *p*_*i*_ are known, the theoretical genetic variance component following Eq. (4) is defined as:

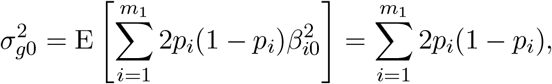

which is a fixed quantity. For a given simulation replicate, we compute the realized variance:

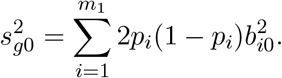

We obtain the desired variance of 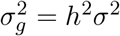 by dividing each *b*_*i*0_ by *s*_*g*0_ (which results in a variance of 1) and then multiply by *hσ*. Combining both steps, the final coefficients *b*_*i*_ used in the algorithm and the corresponding random variable *β*_*i*_ are:

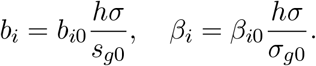

If only 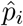 are available, the plug-in Binomial variance estimator 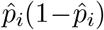 is downwardly biased as shown in Eq. (7). Using that result to unbias our estimator, the initial genetic variance component, which is now a function of two random variables 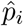 and *β*_*i*0_, is estimated without bias by 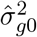, and the realized value of this estimator used in the simulation procedure is denoted 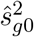 as:

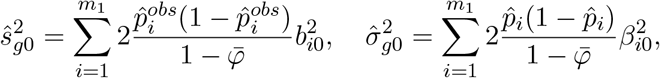

where we denote the realized value of 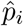 used in simulations as 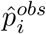 for clarity and 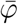 is the mean kinship of Eq. (6). This estimator not only satisfies 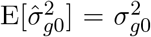, but is a consistent estimator of 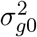 over our random coefficients as the number of independent causal variants *m*_1_ goes to infinity (Lemma 1 in Appendix B). Thus, we define the final coefficients used in a simulation replicate as 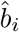, corresponding to the random variable 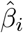 in the model:

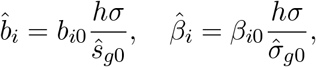

which results in the desired heritability. We obtain reasonable results in practice for large numbers of causal loci (*m*_1_ = 500 in this study) which are not enriched for rare variants, following Theorem 1 in Appendix B.

##### Fixed Effect Sizes (FES) model

In this model, the effect size of locus *i*, defined as 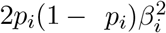, has the same value for every causal locus, so we desire each to equal 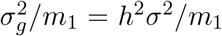. If *p*_*i*_ are known, we simply solve for the desired coefficients:

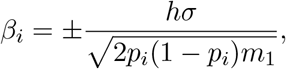

where the signs are chosen randomly with equal probability.

If only 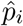 are available, we again unbias the variance estimate following Eq. (7), which results in

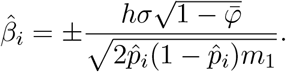

However, unlike RC, the FES estimates are more likely to have biases, meaning that 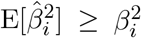 where 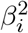 is fixed and non-random in the FES model. This bias may cause a substantial misspecification in heritability estimation, because of the key result in Eq. (8), which follows since the single-locus estimator 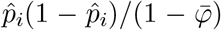 is not consistent as the number of individuals goes to infinity. In particular, this simulation tends to behave poorly if rare variants are causal. As a consequence, the data generated for this paper in this case only used an alternative estimator: let 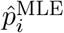 be the maximum likelihood estimate of *p*_*i*_ obtained from the model

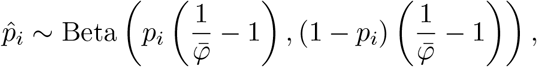

which is the Balding-Nichols model for 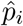 that matches its known moments (Eq. (6)): 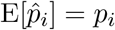 and 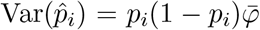. In particular, 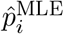 is estimated from 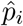 as the only data point, and using the known 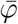 estimate. While this strategy reduces bias somewhat, it does not result in unbiased estimates as shown in the results.

##### Construction of intercept

When *p*_*i*_ are known, we obtain the desired trait mean *µ* by solving for the intercept in Eq. (3), and we use *a* to denote the realized value of *α* used in the simulation:

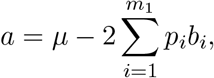

where *b*_*i*_ are computed from the RC or FES procedure above.

If only 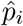 are available, we construct the intercept coefficient using

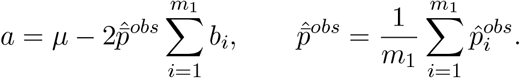

Note that 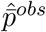 is computed among causal loci only. This works well in practice since in both models E[*β*_*i*_] = 0.

We avoid the naive construction 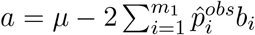, since it is equivalent to centering genotypes at each locus in the model:

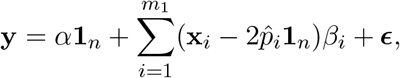

which introduces a distortion in the covariance of the genotypes (Ochoa and Storey, 2021):

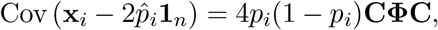

which resembles the standard kinship estimator bias in Eq. (11). These undesirable distortions propagate to the trait, whereas the intercept we constructed does not have these distortions. Our theory proves that such distortions by themselves do not influence heritability estimation using an LMM, since the model’s intercept compensates for this centering (Hou and Ochoa, 2023). Nevertheless, we preserve our trait intercept construction in case future applications are sensitive to the centering effect.

### 2.6 Real data analysis

We utilize the high-coverage NYGC version of the 1000 Genomes Project (Fairley et al., 2020), which is publicly available at ftp://ftp.1000genomes.ebi.ac.uk/vol1/ftp/data_collections/1000G_2504_high_coverage/working/20190425_NYGC_GATK/. We retain only autosomal biallelic SNP loci marked with the filter “PASS”. The final dataset consists of *m* = 91, 784, 660 loci and *n* = 2, 504 individuals. We simulate traits with causal variants that satisfy MAF thresholds in {0, 0.01, 0.05}, and separately, estimate the kinship matrix using genotypes filtered by MAF thresholds in {0, 0.01, 0.05} as well. Mean kinship was estimated by popkin for each MAF and used to simulate traits with matching MAF filter.

The Hispanic Community Health Study / Study of Latinos (HCHS/SOL) is a multi-center study in Hispanic/Latino populations recruited through four centers in Miami, San Diego, Chicago, and the Bronx New York (Sorlie et al., 2010). We used the Phase Ia data available on dbGaP (accession phs000810.v2.p2), which genotyped individuals at 2.5 million SNPs from the Illumina SOL HCHS Custom 15041502 B3 array (core set of SNPs from the Illumina HumanOmni2.5-8 array, with the addition of 110k custom SNPs). We filter genotypes using MAF ≥ 0.01 and HWE ≤ 1e-10. The final dataset consists of *m* = 1, 656, 020 loci and *n* = 11, 721 individuals. We adjust for age and sex in the heritability estimate. Again, we log-transformed all traits except height, % Neutrophils, % Lymphocytes, % Monocytes, % Eosiniphils, % Basophils, and Red Blood Count to improve the model fits.

The San Antonio Family Study (SAMAFS) is a pedigree-based study designed to identify low frequency or rare variants influencing susceptibility to T2D, conducted in 20 Mexican American T2D-enriched pedigrees from San Antonio, Texas (Mitchell et al., 1996). We obtained the exome chip data from dbGaP (accession phs000847.v2.p1). We exclude all unplaced and non-autosomal variants, and further filter using MAF ≥ 0.01 and Hardy-Weinberg Equilibrium p-value (HWE) ≤ 1e-10. The final dataset consists of *m* = 36, 293 loci and *n* = 914 individuals. We also include pedigree information for the estimation of kinship matrix for this data set. We adjust for age and sex in the heritability estimate. Most continuous traits are skewed, so we log-transformed all traits, except height, to improve the model’s fit.

The Nephrotic Syndrome (NS) multiethnic cohort is a multi-ancestry study exploring the etiology of nephrotic syndrome. We analyzed the imputed version of the genotypes filtered using minor allele count ≥ 20, as described elsewhere. The final dataset consists of *m* = 16, 605, 628 loci and *n* = 1, 981 individuals (the 1000 Genomes controls were excluded). We adjust for sex in the heritability estimate. Liability-scale heritability estimates were calculated using the formula of Lee et al. (2011) assuming a prevalence for NS of *K* = 1.69e-4 (Gbadegesin et al., 2015; Jia et al., 2020; Barry et al., 2023), and that SSNS and SRNS are 80% and 20% of NS cases, respectively (Gbadegesin et al., 2015; Jia et al., 2018).

## 3 Results

In order to evaluate the effect of kinship estimation bias on heritability estimation, we consider simulations of genotypes and traits, real genotypes with simulated traits, and lastly real genotypes and traits. Our theory identifies two parameters as key determinants of heritability estimation bias using standard kinship estimators: the mean kinship value 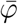, and the proportion of rare variants (in this work defined as the proportion of loci with MAF ≤ 0.05, following Biddanda et al. (2020)). We focused on datasets with higher values of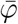, since standard kinship estimators are more biased when there is population structure. On the other hand, the proportion of rare variants in the data, from which kinship estimates are obtained, is highly variable and depends on genotyping platform and quality control filters used, so we consider datasets that span this range (Table 1).

### 3.1 Evaluations based on admixture and family simulations

We begin by considering simulated genotypes and phenotypes where the true heritability is known. The genotype simulations are based on an admixture model, with or without family structure, with parameters chosen previously to match those of real multiethnic datasets (Yao and Ochoa, 2022). The mean kinship parameters of these two simulations are 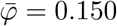 and 0.163 (first replicate, varies slightly per replicate), respectively, which are only slightly larger than those of the real datasets we analyze in this work (1).

First we consider the admixture simulation without family structure. The true kinship matrix of this simulation, as well as estimates for all methods from the first replicate are visualized in Fig. S1. As expected from our theoretical results, only the Popkin kinship estimator produces unbiased heritability estimates, whose distribution closely resembles estimates obtained using the true kinship matrix of the simulation (Fig. 2A and Fig. S2A using traits simulated from the FES and RC models, respectively). The Standard ROM kinship estimator results in a small but noticeable downward bias, in accordance to our theoretical predictions (Fig. 1). Surprisingly, the Standard MOR, which is the most commonly used kinship estimator in practice, but whose bias does not have a closed form, results in considerably more downwardly biased heritability estimates than the ROM counterpart. Furthermore, downward biases are greater for higher true heritability values. To further illustrate the pattern of these biases, we plotted the bias trends for each kinship estimators (Fig. 2B and Fig. S2B).

**Figure 2:**
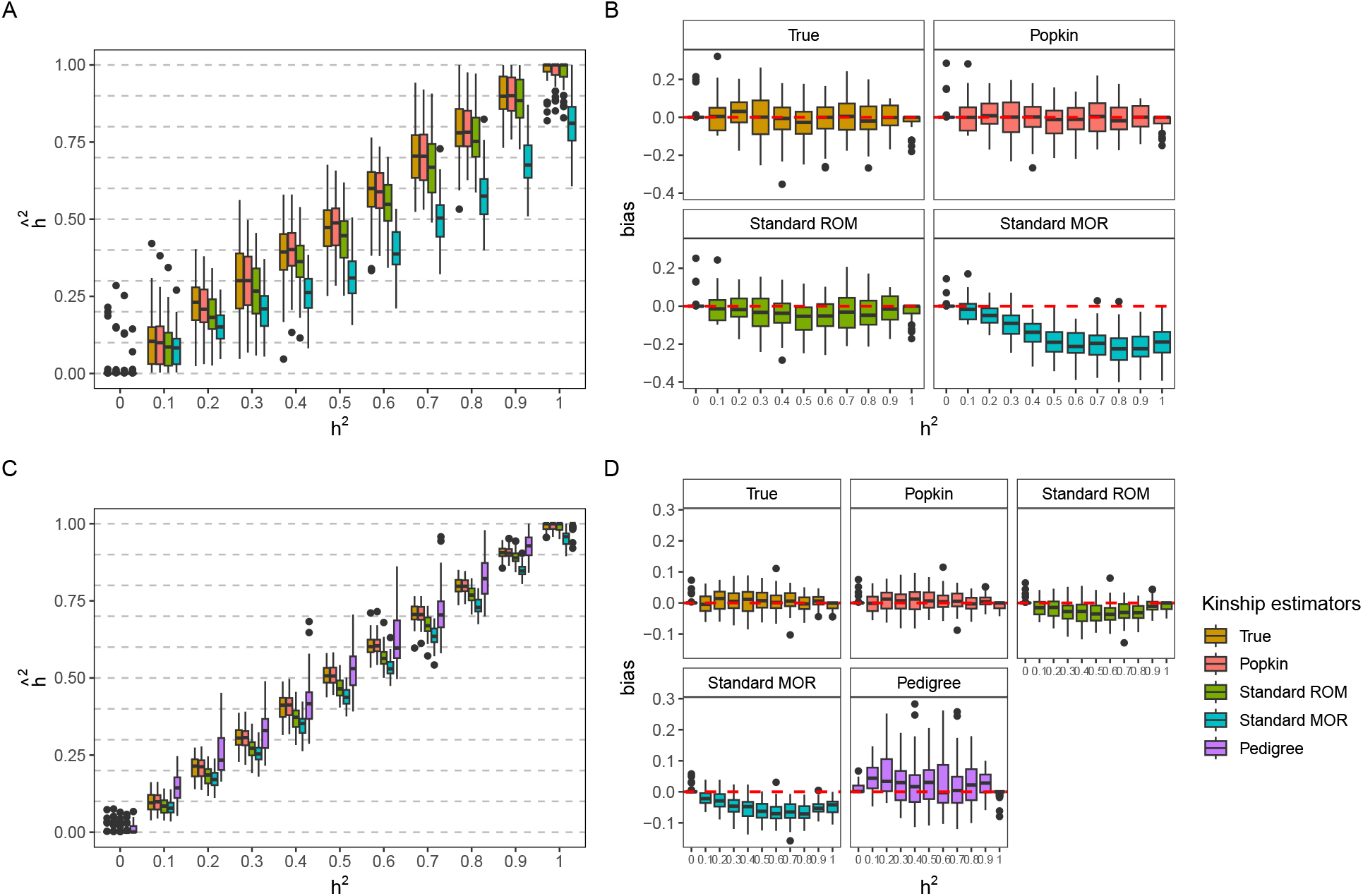
Heritability estimation using GCTA with various kinship estimators evaluated using simulated genotypes and traits. The RC trait model was used to simulate traits in this figure. True heritability values *h*^2^ (x-axis) are compared to their estimates 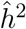 or their bias 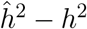. **A-B**. Admixture simulation. **C-D**. Admixture plus 20 generations of family structure simulation. Every true heritability simulation was replicated 50 times.

Next, we consider the simulations with admixture followed by 20 generations of family structures (kinship matrices in Fig. S3). Heritability estimation biases are similar in magnitude in this scenario, but estimation variance is much reduced, apparently a consequence of the presence of closely related individuals. Importantly, the observed trends remain consistent with the previous scenario, namely that Popkin yields unbiased estimates in agreement with the true kinship matrix, while Standard ROM has a consistent downward estimation bias and Standard MOR has an even more severe bias that is higher for larger values of the true heritability (Fig. 2C-D and Fig. S2C-D). Additionally, here we tested using the kinship matrix derived from the true pedigree, which models the relatedness due to family structure but ignores the population structure due to shared ancestry that is also present. Unlike other cases, we find that these pedigree-based estimates are upwardly biased, and also have a much greater variance than the rest of the estimates.

### 3.2 Evaluations based on 1000 Genomes genotypes and simulated traits

The previous simulations differ from real data in their relative dearth of rare variants (MAF ≤ 0.05). In our simulations, ancestral allele frequencies are drawn uniformly (since the maximum MAF is 1/2, the expected proportion of rare variants is 0.05 × 2 = 0.10), but after the admixture and family structure simulation variants tend to become rarer, resulting in a proportion of rare variants of 0.18-0.19 (Table 1). However, in WGS datasets like 1000 Genomes rare variants are the majority (0.93 when there are no MAF filters; Table 1). In order to characterize the effects of rare variants, we used 1000 Genomes genotypes with simulated traits, in this case with a fixed heritability of *h*^2^ = 0.8. 1000 Genomes is a multiethnic cohort which includes African, European, South Asian, East Asian, and Hispanic individuals (kinship matrices in Fig. S4), with a high mean kinship estimate of 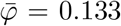 without MAF filter (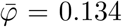 if only including MAF ≥ 0.01, and 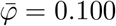 for MAF ≥ 0.05). Of note, in this evaluation we apply MAF filters separately for causal variants (which influence the trait architecture) and for inclusion in the kinship estimates (which influence kinship and downstream heritability estimation biases).

Here we find different behaviors for the two trait models we used, which differ in how they assign coefficients to rare variants. The RC model gave results most consistent with our previous simulations, presented to further solidify Popkin as the only unbiased method, while FES identifies a new source of bias for Standard MOR with some caveats specific to this trait simulation strategy as applied to real genotypes.

The RC (random coefficients) model draws causal coefficients independently of allele frequency, resulting in smaller effect sizes (product of coefficient and genotype variance) for rare variants. Importantly, RC is also the only model guaranteed to result in correctly specified heritabilities when true allele frequencies are unknown, as is the case for real genotypes. Popkin again yields unbiased estimates in most of these cases (overlap with the true value of *h*^2^ = 0.8), although there may be small downward biases if only common variants are causal but rare variants are included in the kinship estimate (Fig. S5). The Standard ROM estimator has a small downward bias in all cases, and always has smaller estimates than Popkin, as predicted by Eq. (12). Lastly, the Standard MOR estimator has its most extreme underestimates of heritability when its kinship estimate includes rare variants, regardless of the presence or absence of rare variants among causal loci. When estimated using only common variants, Standard MOR results in practically unbiased heritability estimates. Thus, these results are consistent with our previous simulations for common variants, but clarify the key role rare variants play in modulating the downward heritability bias resulting from using the Standard MOR estimator.

Next, we turn to traits simulated with the FES (fixed effects sizes) model, which assign causal coefficients that are roughly inverse proportional to the square root of the minor allele frequency (see Methods), thus upweighing rare variants. When causal variants are more common (MAF ≥ 0.01) we saw results as before, namely that Popkin gives relatively unbiased estimates, while Standard MOR is downwardly biased unless only common variants are used to estimate kinship. Unfortunately, this simulation model results in a misspecified heritability when using real genotypes and rare causal variants, since their inverse genotype variance of individual variants cannot currently be estimated without bias (Eq. (8)). Thus, when traits include rare causal variants, heritability estimates were systematically lower than the desired *h*^2^ = 0.8 for all kinship estimators (first column of Fig. 3). Our theory suggests that these trait MAF ≥ 0 results are a bias in how traits are simulated, although at present we cannot disentangle it from estimation biases. Nevertheless, keeping this heritability misspecification in mind, these results yield a new pattern: Standard MOR can yield estimates that exceed those of Popkin and Standard ROM. Although not consistently, we see higher estimates for Standard MOR when the trait has rare causal variants: either no MAF filter for trait and no MAF filter for kinship estimator, no MAF filter for trait and MAF ≥ 0.05 for kinship estimator, and MAF ≥ 0.01 for trait and MAF ≥ 0.05 for kinship estimator. Thus, Standard MOR can result in both downward and upward heritability biases depending on the rare variant architecture of the trait and the inclusion of rare variants in the kinship estimate.

**Figure 3:**
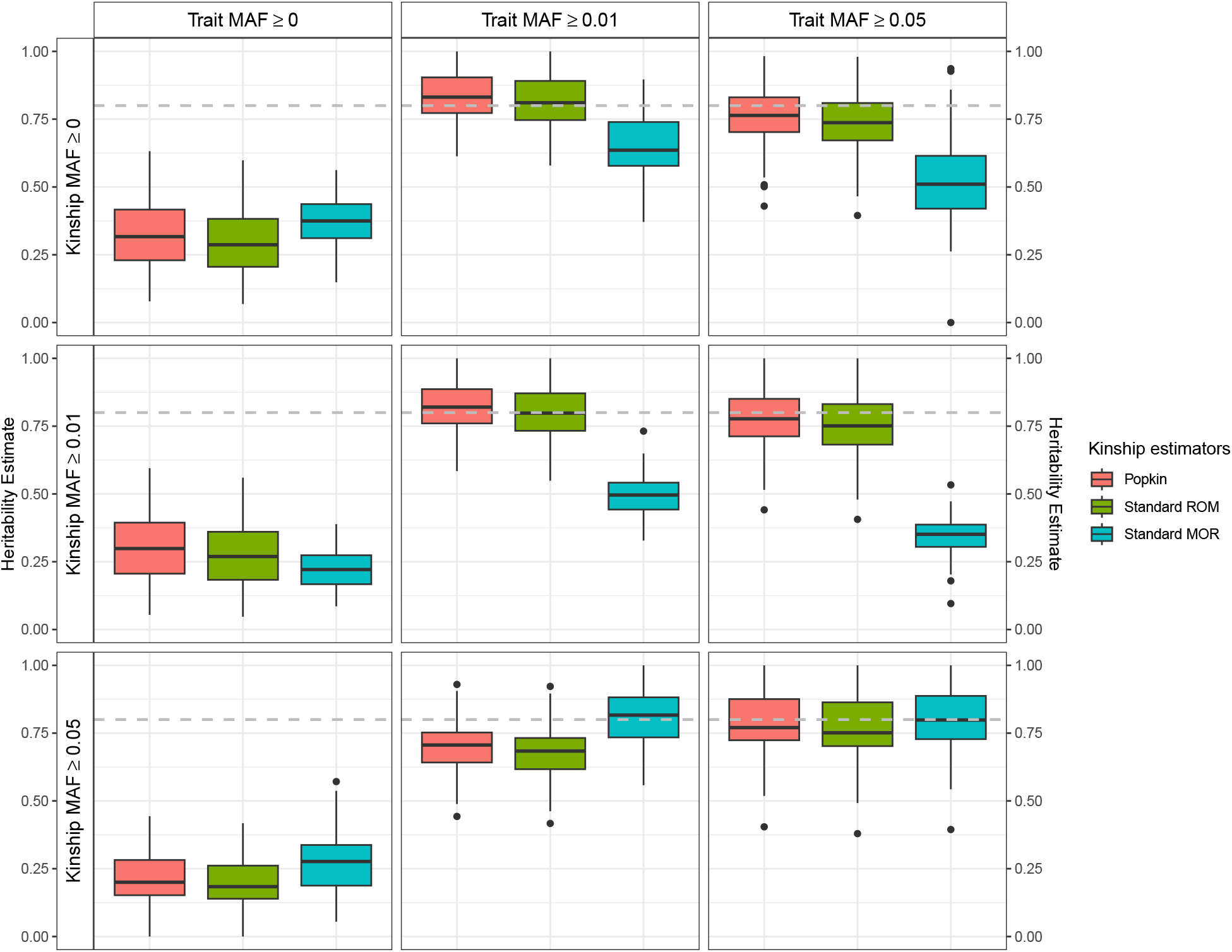
Testing effect of rare variants using 1000 Genomes genotypes and simulated traits. Here we used the FES trait model, which upweighs rare variants and results in unique cases where Standard MOR estimates have upward heritability biases, but has a mispecified heritability for real genotypes and rare caual variants (affects first column only; see Methods). True heritability is *h*^2^ = 0.8 in all panels. Every trait simulation was replicated 50 times. For simulated traits, MAF thresholds shown are applied before causal variants are selected, thus influencing the architecture of the trait. For kinship estimates, MAF thresholds are applied to genotypes before kinship matrices are estimated from these genotypes, thus influencing only kinship estimation.

### 3.3 Analysis of real genotype and phenotype datasets

Having established that only Popkin results in unbiased heritability estimates, and having characterized the biases of the Standard ROM and MOR estimators, we now demonstrate how these biases manifest in real genotype and phenotype data. Recall that the Standard MOR kinship estimator is the most commonly used, being the default for GCTA and related methods. We analyzed three real datasets with complex population structures and otherwise complementary features: the Hispanic Community Health Study/Study of Latinos (HCHS/SOL), the San Antonio Mexican American Family Study (SAMAFS), and the Nephrotic Syndrome (NS) multiethnic cohort.

HCHS/SOL is a population study with strong population structure due to the admixture structure of the Hispanic individuals sampled from various nationalities in this study (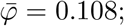Fig. S6). Throughout the 33 traits that we analyzed, Popkin always estimates larger heritabilities than Standard ROM (Fig. 4), as predicted by our theoretical results. These differences are large relative to the standard errors of these estimates, where often the Popkin estimate is about a standard error larger than the Standard ROM estimate. In contrast, Standard MOR estimates are often the largest of the three estimates, again by up to a standard error, although for some traits they are instead smaller or similar to the Popkin estimates. The proportion of rare variants, which influences the kinship estimates and the bias of Standard MOR in particular, is the lowest among the real datasets (0.26), but it is slightly higher than the simulated data (Table 1). Thus, we do not expect the downward biases due to rare variants in the Standard MOR kinship estimate, while a large proportion of rare causal variants (which are likely not genotyped) likely explains the upward bias of Standard MOR in most of these traits. If so, traits with larger Standard MOR versus ROM estimates may have more rare causal variants than traits otherwise. Interestingly, the estimated heritability of height is high for all estimators, just under the *h*^2^ = 0.8 estimated from classic sibling studies, which is achieved here only after conditioning for sex and age as fixed effects.

**Figure 4:**
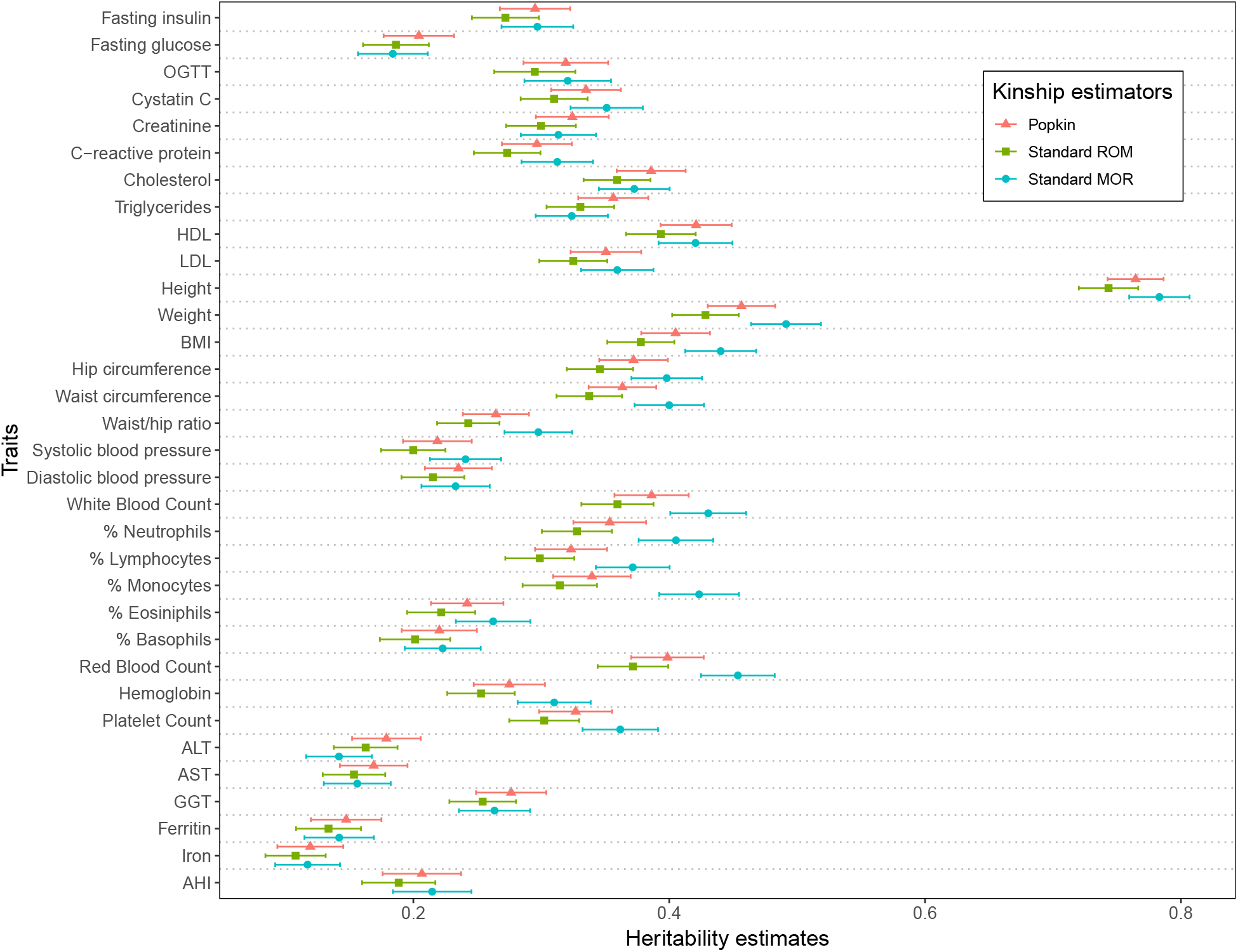
Heritability estimates on Hispanic Community Health Study/Study of Latinos (HCHS/SOL). The figure compares heritability estimates with GCTA using Popkin, Standard ROM, and Standard MOR kinship estimators.

The SAMAFS data is considerably less structured, potentially since it is a more homogeneous set of Hispanics sampled from a single city and belonging to only 20 families (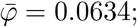Fig. S7). Although we see the same patterns as in HCHS/SOL, namely that Popkin yields larger estimates than Standard ROM, differences are much smaller here relative to their standard errors (Fig. 5). Interestingly, unlike HCHS/SOL, here Standard MOR and ROM produce very similar estimates, which could again be partly due to a low proportion of rare variants used to estimate kinship (0.28; Table 1), although upward biases due to causal rare variants were not observed either. Furthermore, SAMAFS has a well-documented pedigree structure, which we use to estimate kinship to mimic family analyses, although it does not model the population structure due to admixture expected for Hispanics. Pedigree heritability estimates tended to be larger, sometimes considerably so relative to the standard error (for example, Adiponectin), whereas for one trait it is the smallest estimate (Fasting glucose). Since pedigree-based heritability estimates were upwardly biased and highly variable in our simulations (Fig. 2), we conclude that these estimates are likely also upwardly biased. We again see high height heritability estimates, also obtained by conditioning for sex and age, which is less surprising for a family study.

**Figure 5:**
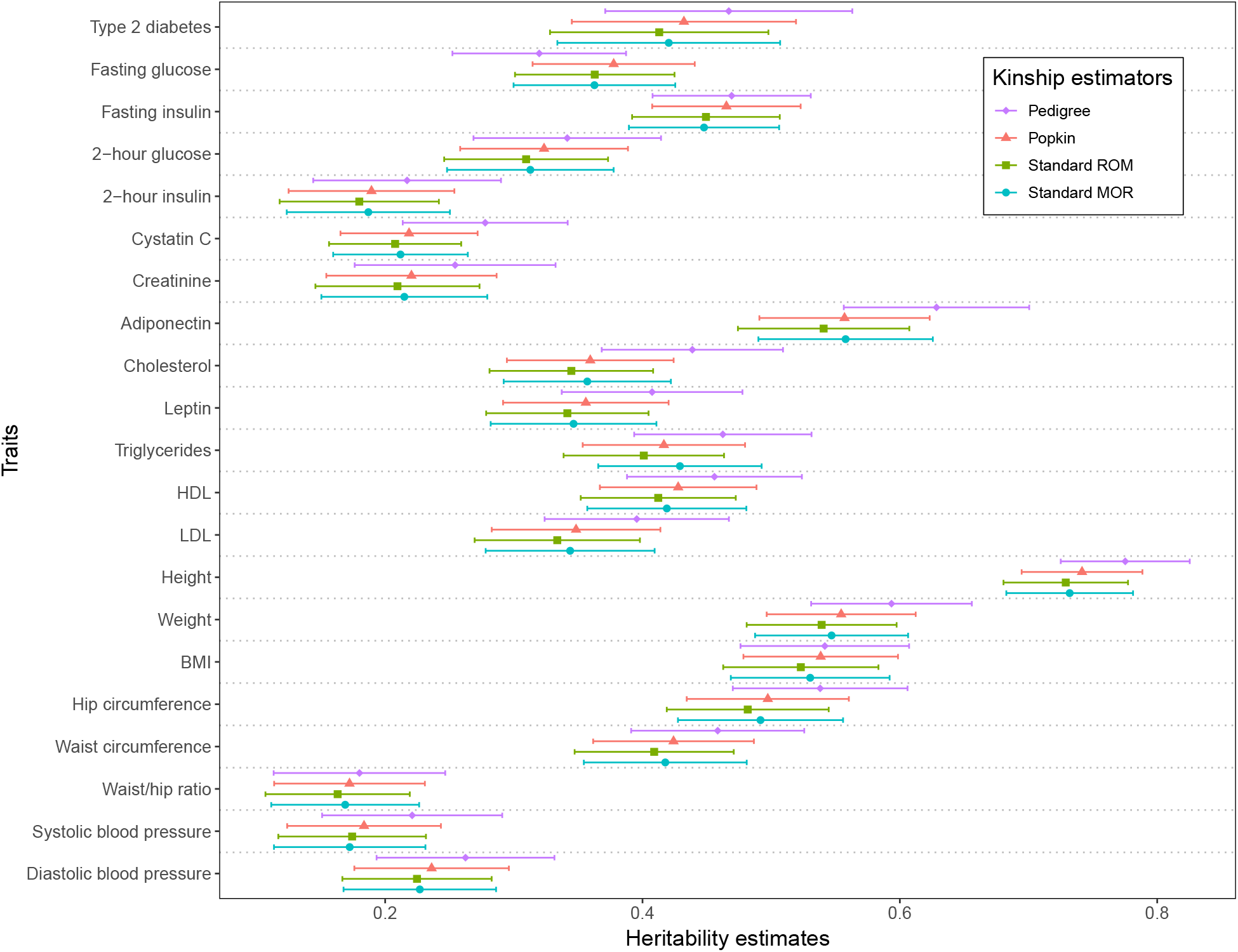
Heritability estimates on San Antonio Mexican American Family Study: Type 2 Diabetes (SAMAFS). Heritability estimates are calculated using GCTA and the Pedigree, Popkin, Standard ROM, and Standard MOR kinship estimates.

Since HCHS/SOL and SAMAFS are studies of Hispanics that overlap on 16 traits, we compared their heritability estimates. Despite the potential for heritability differences due to differences in environment or genetics, we find high Pearson correlations of ≈ 0.8 between the estimates of each method in both datasets (Fig. S8). We also observe slightly higher heritability estimates in SAMAFS compared to HCHS/SOL. However, for this small number of traits there are no clear differences in consistency between Popkin, Standard ROM and MOR estimates, whose regression lines all closely follow the *y* = *x* line.

The NS dataset is a multiethnic dataset including large proportions of African, European, South Asian, and admixed ancestry individuals (Fig. S9). Its mean kinship value of 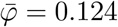 is the largest analyzed with real phenotypes, and just below that of 1000 Genomes. This dataset has three binary traits: each of NS and its subtypes SSNS (steroid sensitive NS) and SRNS (steroid resistant NS) contrasts individuals with those diseases to control samples. As before, Popkin estimates are larger than those of Standard ROM, although the differences are negligible. However, here Standard MOR yields considerably smaller heritability estimates than Popkin, a pattern that was not observed in the other datasets (Fig. 6). The proportion of rare variants is second highest here (0.56; Table 1), which likely explains the severe downward biases observed for Standard MOR. The visualized estimates are in the liability scale. We noticed that Popkin estimates in the observed scale were the maximum possible (0.999) for NS and SSNS (Table S1), yet liability scale estimates are less than 0.3, so we suggest interpreting these estimates with caution.

**Figure 6:**
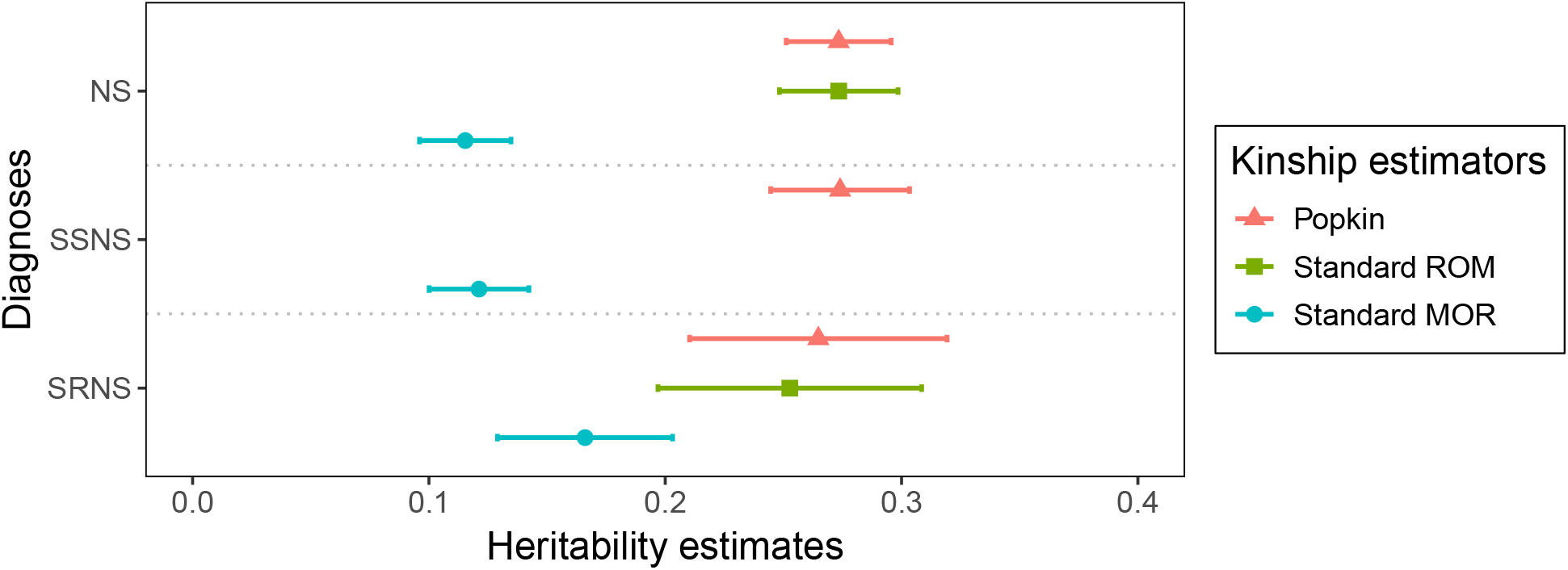
Heritability estimates on Nephrotic Syndrome (NS) dataset. The figure shows heritability estimates in the liability scale using Popkin ROM, Standard ROM, and Standard MOR kinship estimators on the NS multiethnic cohort. The binary traits are NS (NS vs control), SSNS (steroid sensitive NS subset vs control), and SRNS (steroid resistant NS subset vs control). Note the Standard ROM result for SSNS did not converge. For observed scale estimates, see Table S1.

## 4 Discussion

Previous research has shown that commonly used kinship estimators are biased when there is population structure (Ochoa and Storey, 2021). These kinship estimators are widely used to conduct association studies, directly in linear mixed-effects models or indirectly via PCA, models that are remarkably robust to these kinship estimation biases (Hou and Ochoa, 2023). In this study, we characterized the effect of kinship estimation bias on heritability estimation using variance components models, finding that this bias carries over to heritability estimation, in agreement with recent work (Chen and Storey, 2022). Our findings are summarized in Fig. 7. Specifically, when there is population structure, he Standard ROM kinship estimator systematically underestimates heritability, with a bias that has a closed form and depends on the mean kinship value. The much more common Standard MOR estimator does not have a closed form limit, but empirically it has even more severe biases than ROM, which depend on the presence and role of rare variants. Kinship estimated from pedigrees appear to result in upward heritability biases and high variance when there is population structure. Lastly, the Popkin estimator resulted in unbiased heritability estimates.

**Figure 7:**
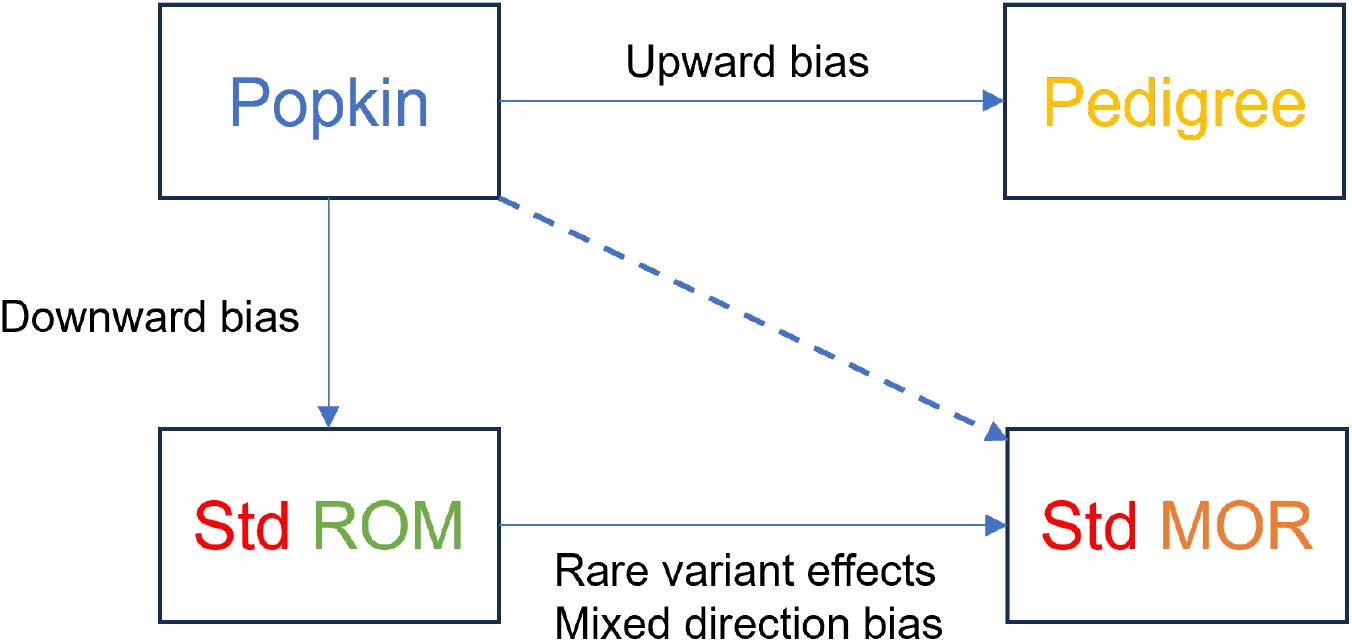
Relationships between kinship estimators in terms of their effects on heritability estimation. Under population structure, Popkin is generally unbiased, Standard ROM is consistent downwardly biased, and pedigree-based kinship estimates are upwardly biased. However, the more common Standard MOR estimator can have either downward or upward biases, depending on whether rare variants are used to estimate kinship or traits have rare causal variants, respectively.

Rare variants can affect heritability estimation in two ways: by introducing biases in kinship estimation, and by having large effects as causal variants. Our evaluations demonstrate that the Standard MOR estimator has stronger biases than the Standard ROM estimator, but the direction of the bias depends on the properties of the rare variant effects. In particular, when rare variants are not causal or when they have small effect sizes, Standard MOR is more downwardly biased than Standard ROM, which is itself always downwardly biased. In contrast, when there is a large proportion of rare causal variants and they have large effect sizes, the Standard MOR has an upward bias. Our hypothesis is that this behavior is due to Standard MOR upweighing rare variants compared to the other two estimators. In HCHS/SOL, many traits appear to have an architecture where rare variants have significant effects, since Popkin’s heritability estimates are lower than those of Standard MOR. However, it remains an open challenge to quantitatively determine the direction and magnitude of the heritability estimation bias due to using the Standard MOR kinship estimator.

Although we drew firm conclusions about the effects of rare variants on heritability estimation across kinship estimators, our work has limitations in how rare variants and their effect sizes are simulated. Our admixture and family simulations begin from pre-existing ancestral variation and do not introduce new mutations, so they primarily generate common variants. To simulate traits influenced by more realistic rare variants, we utilized the genotypes from the 1000 Genomes Project. When rare variants were causal under the FES model, all kinship estimators underestimated heritability (Fig. 3), whereas this bias was not present under the RC model (Fig. S5). This discrepancy reflects a limitation of the FES trait simulation algorithm when applied to real genotypes, for which true ancestral allele frequencies are unknown, leading to upwardly biased estimates of the inverse of genotype variance and misspecified heritability. Overall, estimation of heritability due to rare variants remains a theoretically underdeveloped area, necessitating further research.

Our model has another limitation: it assumes that rare and common variants have the same covariance structure, although there is evidence to the contrary (Zaidi and Mathieson, 2020). In our 1000 Genomes simulation we applied different MAF thresholds to causal variants and to kinship estimation, allowing mismatches in terms of rare variant content between the estimated kinship matrix and the true trait covariance. Popkin is clearly unbiased when causal variants are drawn from the same set used to estimate kinship (diagonal cases in Fig. S5), while we observed small biases for mismatched causal variants and kinship matrices (off-diagonal cases in the figure). Thus, our data supports the hypothesis that rare and common variants have somewhat different structures, although the resulting bias due to ignoring their distinction was relatively small compared to the biases resulting from using the more common Standard MOR estimator.

Pedigrees are available for the San Antonio Family Study (SAMAFS), allowing us to estimate heritability using the pedigree in a manner that more closely resembles classic family studies (Almasy and Blangero, 1998b) as well as more recent approaches from large EHR or insurance claim data (Polubriaginof et al., 2018; Wang et al., 2017b). Notably, SAMAFS was originally analyzed using the pedigree to partition phenotypic variance more finely than our present analysis (Mitchell et al., 1996). However, our simulation results indicate that these heritability estimates are upwardly biased when there is population structure, since the kinship estimate is missing the additional relatedness due to shared ancestry. In practice, incomplete pedigrees also result in unmodeled cryptic relatedness. Further, studies based on close relatives may overestimate heritability due to epistasis effects (Hemani et al., 2013; Young and Durbin, 2014). Supporting this, heritability estimates from close relatives are often inflated due to shared environmental effects, further highlighting the limitations of pedigree-based methods (Zaitlen et al., 2013).

In the Nephrotic Syndrome (NS) multiethnic cohort, traits are binary (case vs. control). In our study, although the liability-scale heritability estimates ranged from 0.1 to 0.3 after transformation, the observed heritability estimates for NS and SSNS using the popkin estimator were 0.99 (Table S1), which occurred because 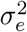 converged to nearly zero. Liability models rely on untestable assumptions (e.g., multivariate normality of latent liability) that, when violated, can produce severely biased estimates (Benchek and Morris, 2013). Our results extend these concerns, suggesting that even when liability-scale estimates appear plausible, they may mask extreme variance components. In addition, it is known that NS, and SSNS in particular, have a single large effect locus in the HLA region, which affects the assumptions of heritability estimation using variance components (Gbadegesin et al., 2015; Adeyemo et al., 2018; Debiec et al., 2018; Jia et al., 2018; Dufek et al., 2019; Jia et al., 2020; Barry et al., 2023). Nevertheless, this data also serves as another illustration of the heritability biases present in the Standard estimators compared to Popkin.

Overall, our study highlights the importance of selecting appropriate kinship estimators for heritability analysis, particularly in structured populations and in the presence of rare variants. These findings provide key insights into the biases inherent in existing kinship estimators and underscore the need for future research to refine estimation methods for more accurate and reliable heritability inference across diverse genetic datasets.

## Acknowledgments

This work was funded in part by the Duke University School of Medicine Whitehead Scholars Program, a gift from the Whitehead Charitable Foundation. Thanks to Rasheed Gbadegesin for sharing his NS data. Thanks to Tiffany Tu for sharing processed data and derivatives for the NS data, and a trait transformation analysis for SAMAFS and HCHS/SOL datasets. Thanks to Grace Rhodes for developing the maximum likelihood estimator of allele frequency used for fixed effect size simulation for real genotype data.

The 1000 Genomes data were generated at the New York Genome Center with funds provided by NHGRI Grant 3UM1HG008901-03S1.

The Hispanic Community Health Study/Study of Latinos is supported by contracts from the National Heart, Lung, and Blood Institute (NHLBI) to the University of North Carolina at Chapel Hill, Chapel Hill, NC (N01-HC65233), University of Miami, Miami, FL (N01-HC65234), Albert Einstein College of Medicine, Bronx NY (N01-HC65235), University of Illinois, Chicago IL (N01-HC65236), and San Diego State University, San Diego CA (N01-HC65237). The following Institutes/Centers/Offices contribute to the HCHS/SOL through a transfer of funds to the NHLBI: National Center on Minority Health and Health Disparities, the National Institute of Deafness and Other Communications Disorders, the National Institute of Dental and Craniofacial Research, the National Institute of Diabetes and Digestive and Kidney Diseases, The National Institute of Neurological Disorders and Stroke, and the Office of Dietary Supplements. The authors thank the staff and participants of the HCHS/SOL study for their important contributions.

The research reported in this article was supported by National Institutes of Health grants. The genetic and phenotypic data were provided by the San Antonio Family Heart Study (SAFHS) investigators and supported by the National Heart, Lung, and Blood Institute (NHLBI) [P01 HL045222] and the San Antonio Family Diabetes/Gallbladder Study (SAFDGS) investigators and supported by the National Institute of Diabetes and Digestive and Kidney Diseases (NIDDK) [R01 DK047482, R01 DK053889]. The phenotypic data were also provided by the Veterans Administration Genetic Epidemiology Study (VAGES) investigators and supported by the Health Services Research and Development, U.S., Department of Veteran Affairs and the Family Investigation Acknowledgement Statement : of Nephropathy and Diabetes (FIND) - San Antonio (FIND-SA) Component and its extension called the Extended FIND (E-FIND) [U01 DK57295] investigators and supported by the NIDDK. The SAFHS gene expression assays were supported by a donation from the Azar and Shepperd families. The exome sequencing, exome chip genotypic, and whole genome sequencing data were provided by the T2D-GENES Consortium grants U01 DK085524, U01 DK085584, U01 DK085501, U01 DK085526, and U01 DK085545 and investigators supported by the NIDDK. This manuscript was not prepared in collaboration with investigators of the T2D-GENES SAMAFS/Consortium and does not necessarily reflect the opinions or views of the members of the T2D-GENES SAMAFS/Consortium, or the NIDDK

## Data and code availability

The data and code generated during this study are available on GitHub at https://github.com/OchoaLab/bias-herit-paper

## Appendices

### A Variance of the plug-in Binomial variance estimator 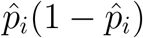

To explain the inconsistency of the FES variance estimator, as well as the Standard MOR kinship estimator, we calculate the variance of 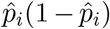 shown in the main Methods, where 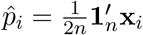 is the empirical allele frequency estimated from genotypes **x**_*i*_.

We begin by rewriting the quantity of interest as a bilinear form:

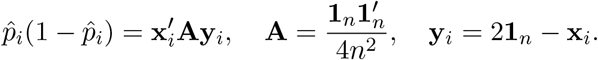

This allows us to use a known identity to complete the variance calculation, which assumes multivariate normal distributions for **x**_*i*_ and **y**_*i*_ and a symmetric **A**, namely

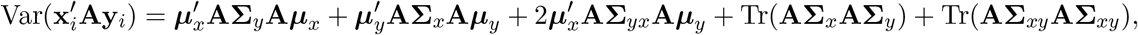

where ***µ***_*x*_, ***µ***_*y*_ are expectation vectors, and **Σ**_*x*_, **Σ**_*y*_, **Σ**_*xy*_ are covariance and cross-covariance matrices, respectively. Thus, we assume a normal version of the kinship model of Eq. (1), namely

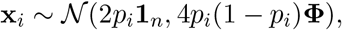

which is expected to result in a more accurate calculation for *p*_*i*_ near 0.5 (high MAF). It follows directly from these assumptions that **y**_*i*_ is also normal and has moments

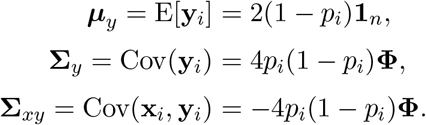

Substituting in, gathering like terms and simplifying, we obtain that:

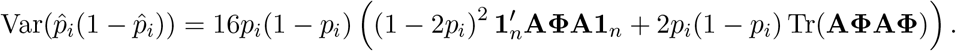

Since 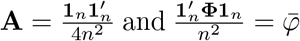, and using the cyclic property of traces, the following terms simplify:

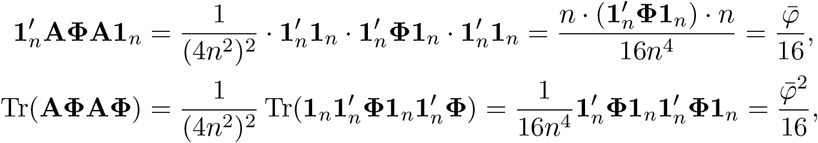

Thus, the final variance equation is:

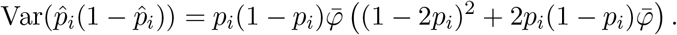

Note that even though 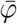 is defined as a sample parameter (for finite *n*), its value will converge to the population value as *n* grows, and this value will be non-zero when there is population structure.

Note that

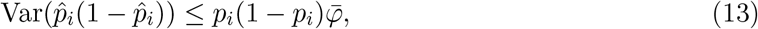

since the second factor 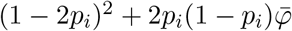 is a quadratic on *p*_*i*_ with values of 1 at both edge points (*p*_*i*_ = 0 and *p*_*i*_ = 1) and a minimum at *p*_*i*_ = 1*/*2 that evaluates to 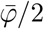, so the whole second factor is non-negative and bounded above by 1. Thus, the inequality is tighter near the edges, and looser in the middle (near *p*_*i*_ = 1*/*2).

### B Consistency of the RC initial variance estimator 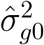 and the final variance 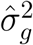

#### Lemma 1.

*Let* 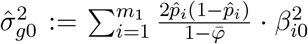, *where β*_*i*0_ ∼ 𝒩 (0, 1) *i*.*i*.*d*., *and* 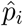 *are allele frequency estimates satisfying:*

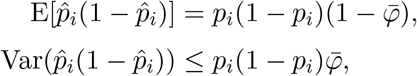

*with fixed p*_*i*_ ∈ [*δ*, 1 − *δ*] *for some δ >* 0, *and each term in the sum is independent across i. Define the target initial genetic variance component as*

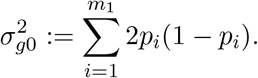

*Then, as m*_1_ → ∞,

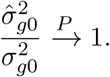

*That is*, 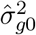 *is a consistent estimator of* 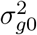.

*Proof*. Define 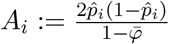, so that 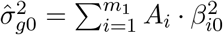. Since *β*_*i*0_ ∼ 𝒩 (0, 1) and are independent of *A*_*i*_, we have:

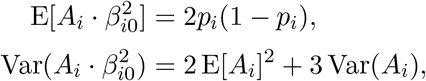

because 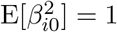 and 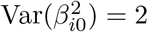. Then 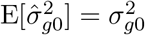.

Therefore:

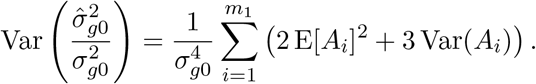

It follows from our assumptions that:

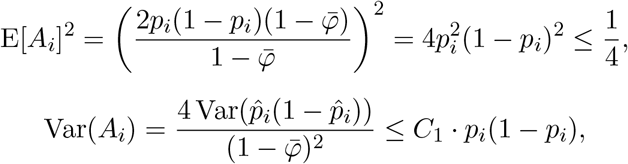

for constants *C*_1_ depending on 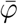.

Thus, for some constant *C*_2_:

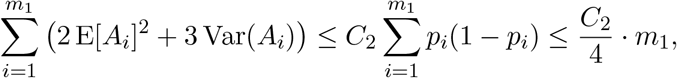

since *p*_*i*_(1 − *p*_*i*_) ≤ 1*/*4 for all *i*, the right-hand side grows at most linearly with *m*_1_, and hence the numerator of the variance expression increases at most proportionally to the number of causal variants *m*_1_.

Under the assumption that all *p*_*i*_ ∈ [*δ*, 1 − *δ*], each term *p*_*i*_(1 − *p*_*i*_) ≥ *δ*(1 − *δ*), so the initial variance satisfies

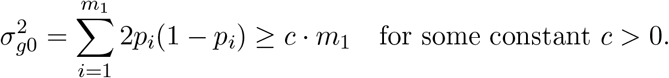

Therefore, as *m*_1_ → ∞:

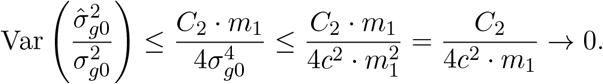

Therefore, by Chebyshev’s inequality,

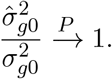

□

**Remark**. When causal variants have small minor allele frequencies (i.e., small *p*_*i*_), the terms *p*_*i*_(1 − *p*_*i*_) become small, reducing the denominator 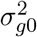. This leads to a looser (i.e., larger) upper bound in the variance of 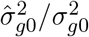, and thus slower convergence in probability.

#### Theorem 1

(Consistency of Final Genetic Variance). *Let* 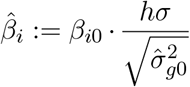 *be the final standardized effect sizes constructed from i*.*i*.*d. draws β*_*i*0_ ∼ 𝒩 (0, 1), *and define the final genetic variance as*

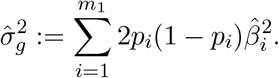

*Let the target variance be* 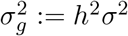. *Then, under the assumptions of Lemma 1*,

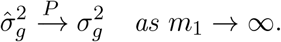

*Proof*. By Lemma 1, we have

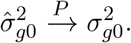

Then,

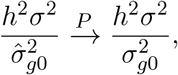

by the continuous mapping theorem.

Substituting into the definition of 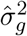, we have:

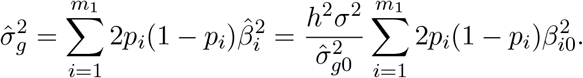

The sum 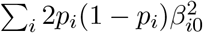 is a consistent estimator of 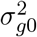 by construction, so

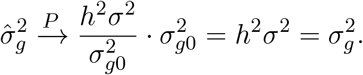

Hence, 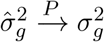, as claimed. □

## S1 Supplemental Tables and Figures

**Table S1:**
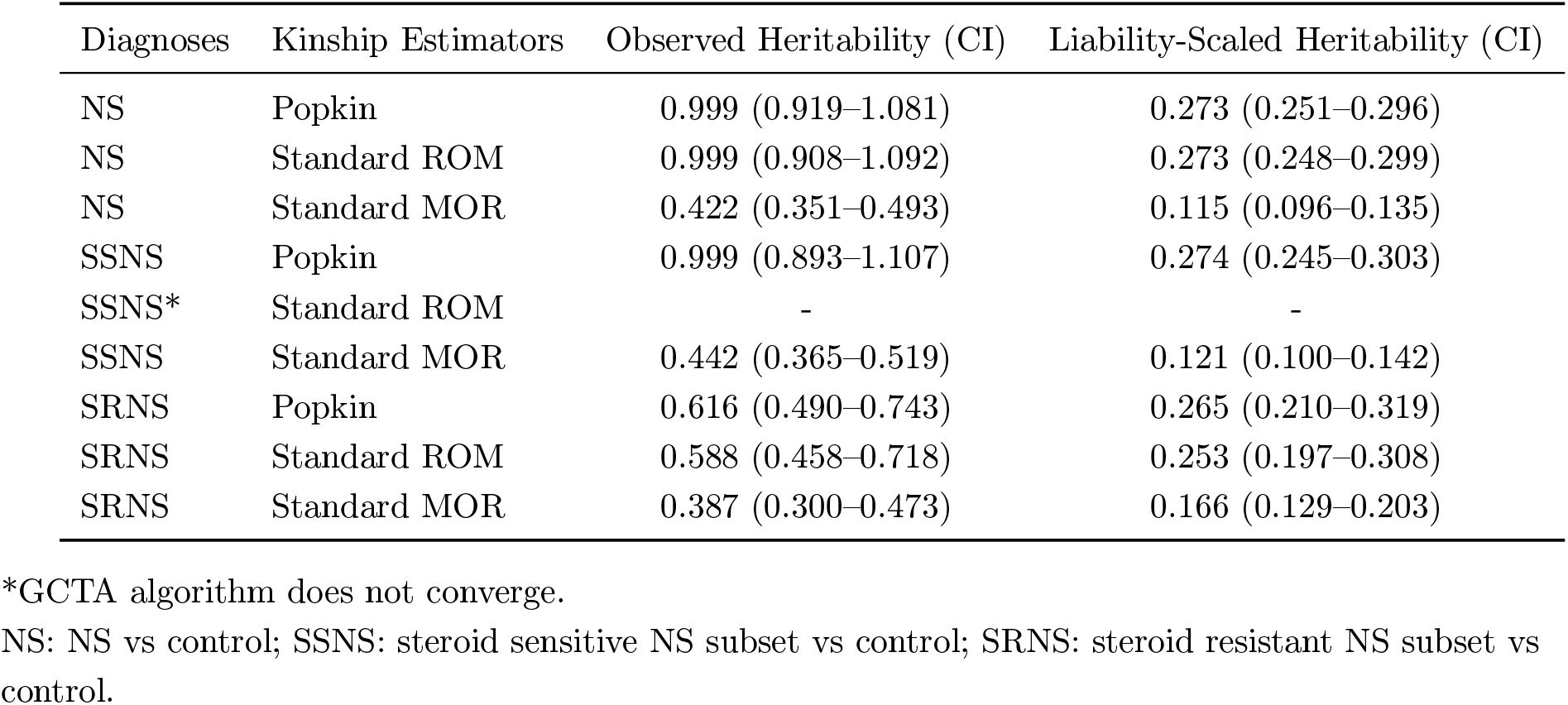
Heritability Estimates for the NS multiethnic cohort.

**Figure S1:**
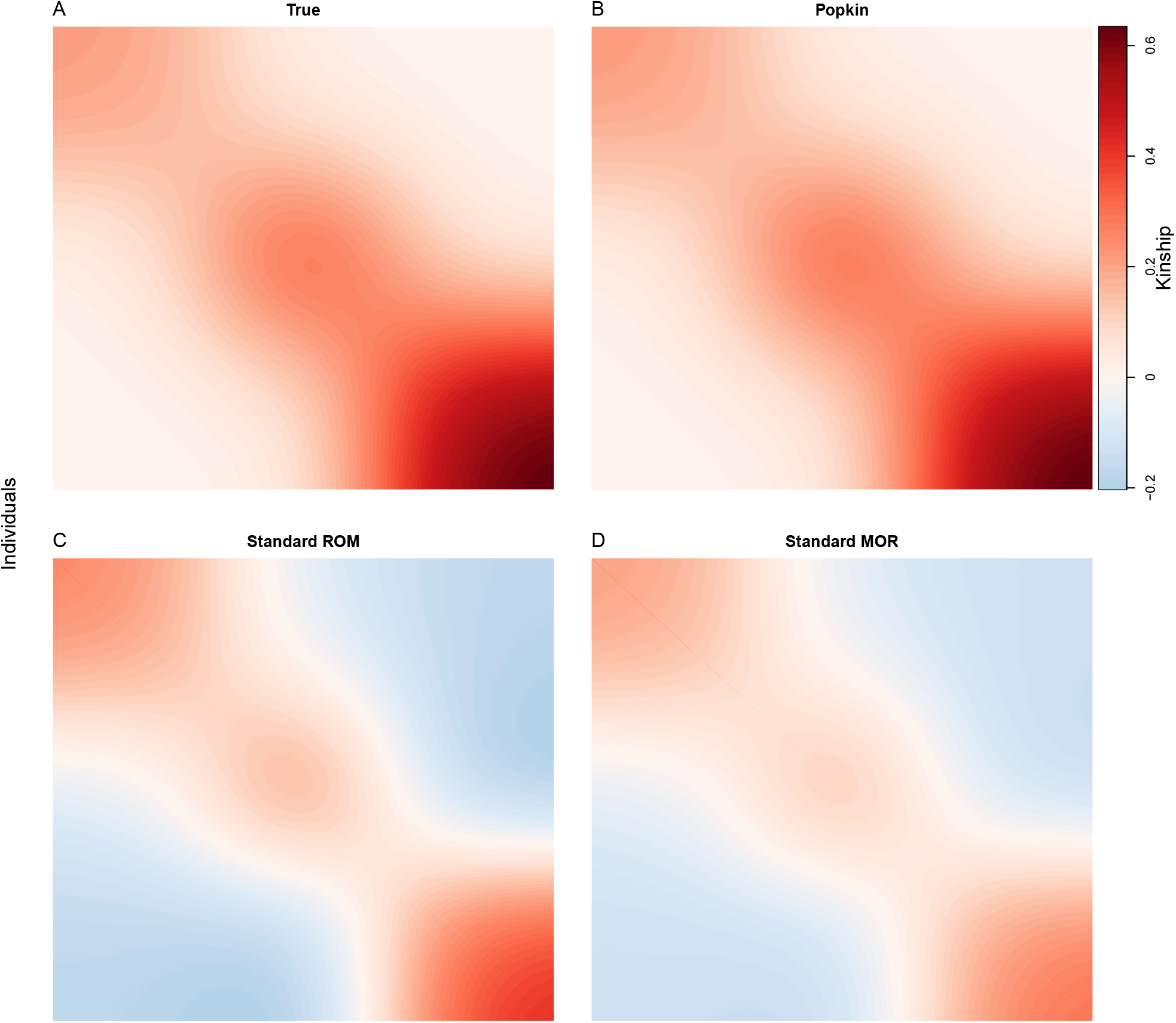
Kinship matrix estimates for the simulated admixture dataset. Estimates are calculated on the genotype matrix of the first replicate of this simulation. In each panel, individuals are placed along both x and y axes, and the kinship value *φ*_*jk*_ of a pair of individuals *j* and *k* is visualized as color, where values near zero appear white, large positive values are darker red, and large negative values are darker blue. The diagonal displays inbreeding values *f*_*j*_ = 2*φ*_*jj*_ − 1 rather than self-kinship values *φ*_*jj*_ because the inbreeding value of an individual equals the kinship of the parents, so it is on the same scale as kinship whereas self-kinship is not. **A**. True kinship matrix of the simulation, calculated from the admixture model parameters as in Ochoa and Storey (2021). **B**. Popkin estimate, which is unbiased. **C**. Standard ROM (ratio-of-means) estimate, which has a bias described in the Methods. **D**. Standard MOR (mean-of-ratios) estimate, which has a similar bias to Standard ROM but has additional biases that do not have a closed form; the difference is driven by rare variants, which are upweighed in the MOR version.

**Figure S2:**
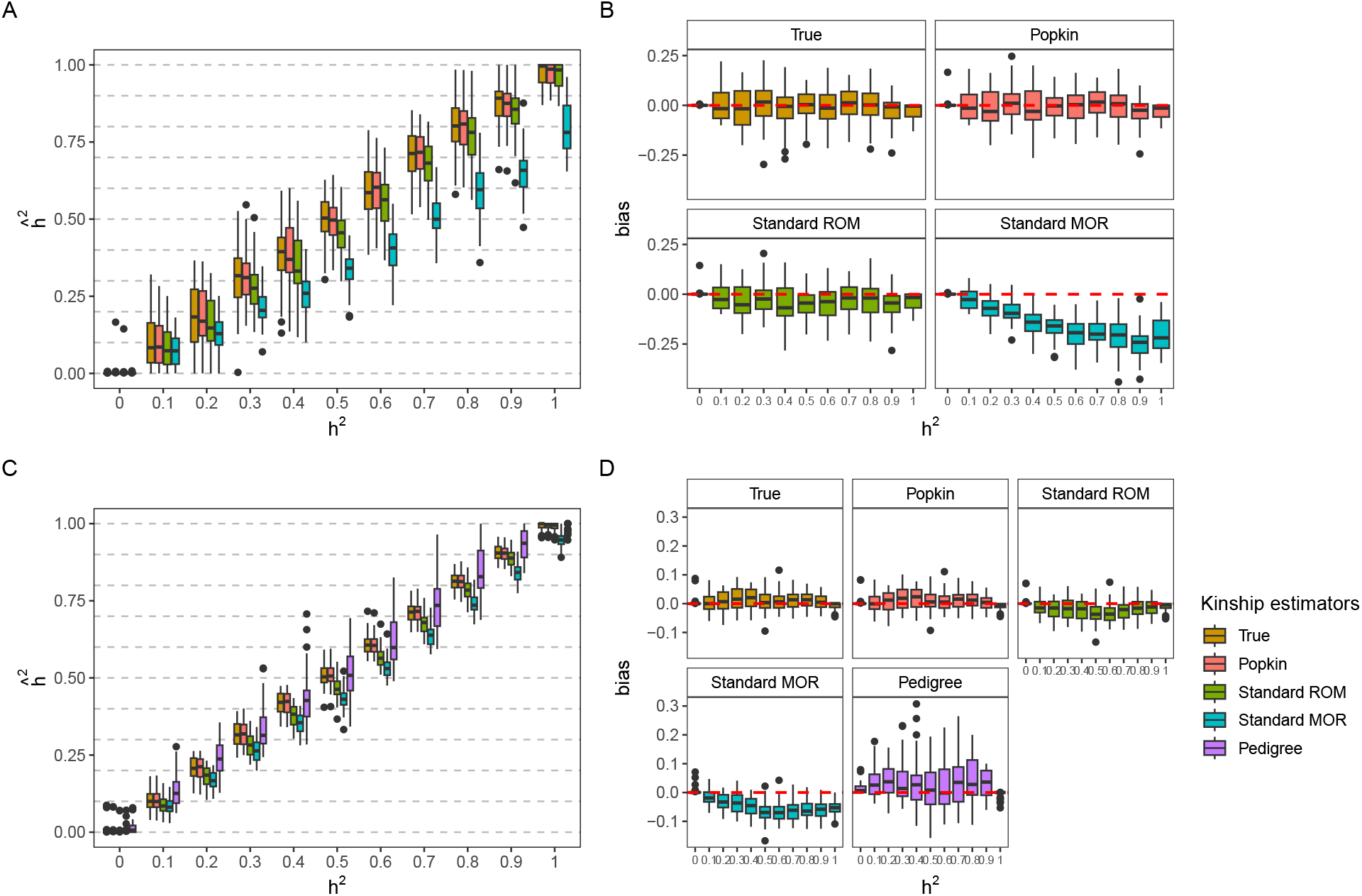
Heritability estimation using GCTA with various kinship estimators evaluated using simulated genotypes and traits. The FES trait model was used to simulate traits in this figure. **A-B**. Admixture simulation. **C-D**. Admixture plus 20 generations of family structure simulation. See Fig. 2 for more information.

**Figure S3:**
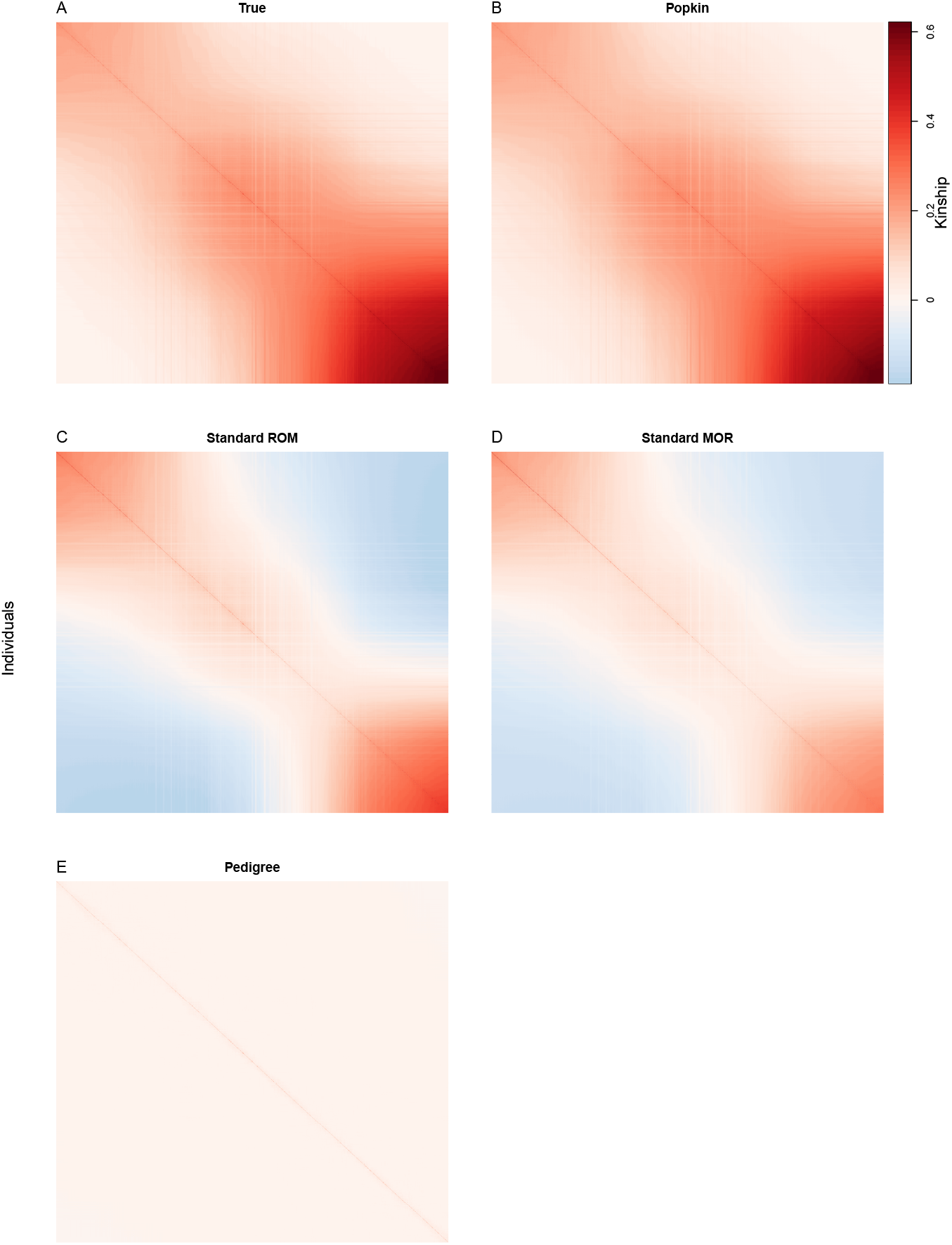
Kinship matrix estimates for the simulated admixture plus family structure dataset. Estimates are calculated on the genotype matrix of the first replicate of this simulation. The simulation reorders individuals so that families appear closer together along the diagonal (notice darker reds corresponding to higher kinship values). Darker or brighter lines in these figures correspond to individuals with more different ancestry than their neighbors on average (including family members such as spouses) which occur with some frequency in this simulation. See Fig. S1 for more details, including descriptions of panels, except: **E**. The kinship matrix calculated from the true pedigree of the simulation using standard methods, which erroneously treat founders as unrelated even though in this simulation they are admixed so their relatedness is as in Fig. S1A.

**Figure S4:**
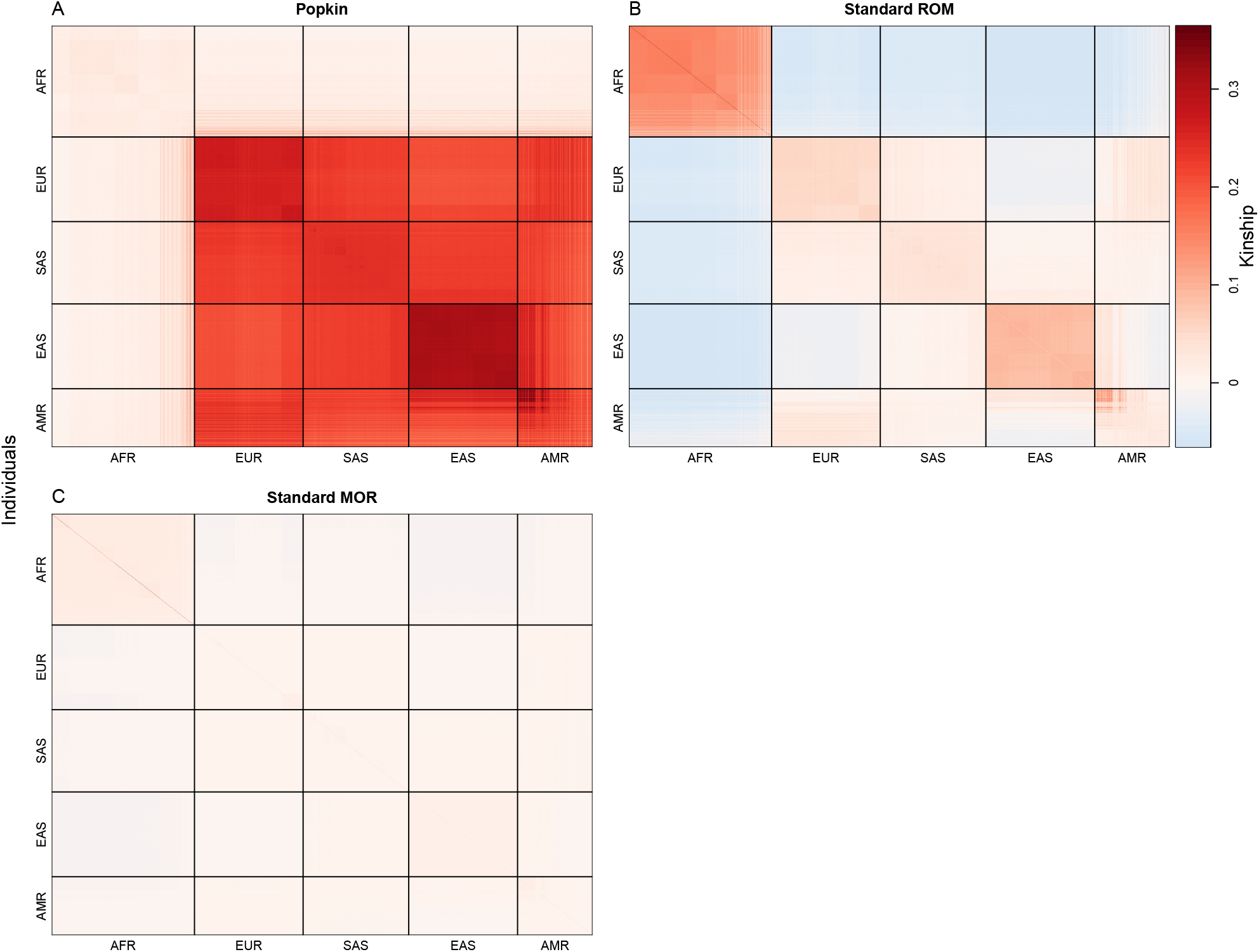
Kinship matrix estimates for the 1000 Genome dataset. AFR = African, EUR = European, SAS = South Asian, EAS = East Asian, AMR = Admixed Americans (Hispanics). See Fig. S1 for more details.

**Figure S5:**
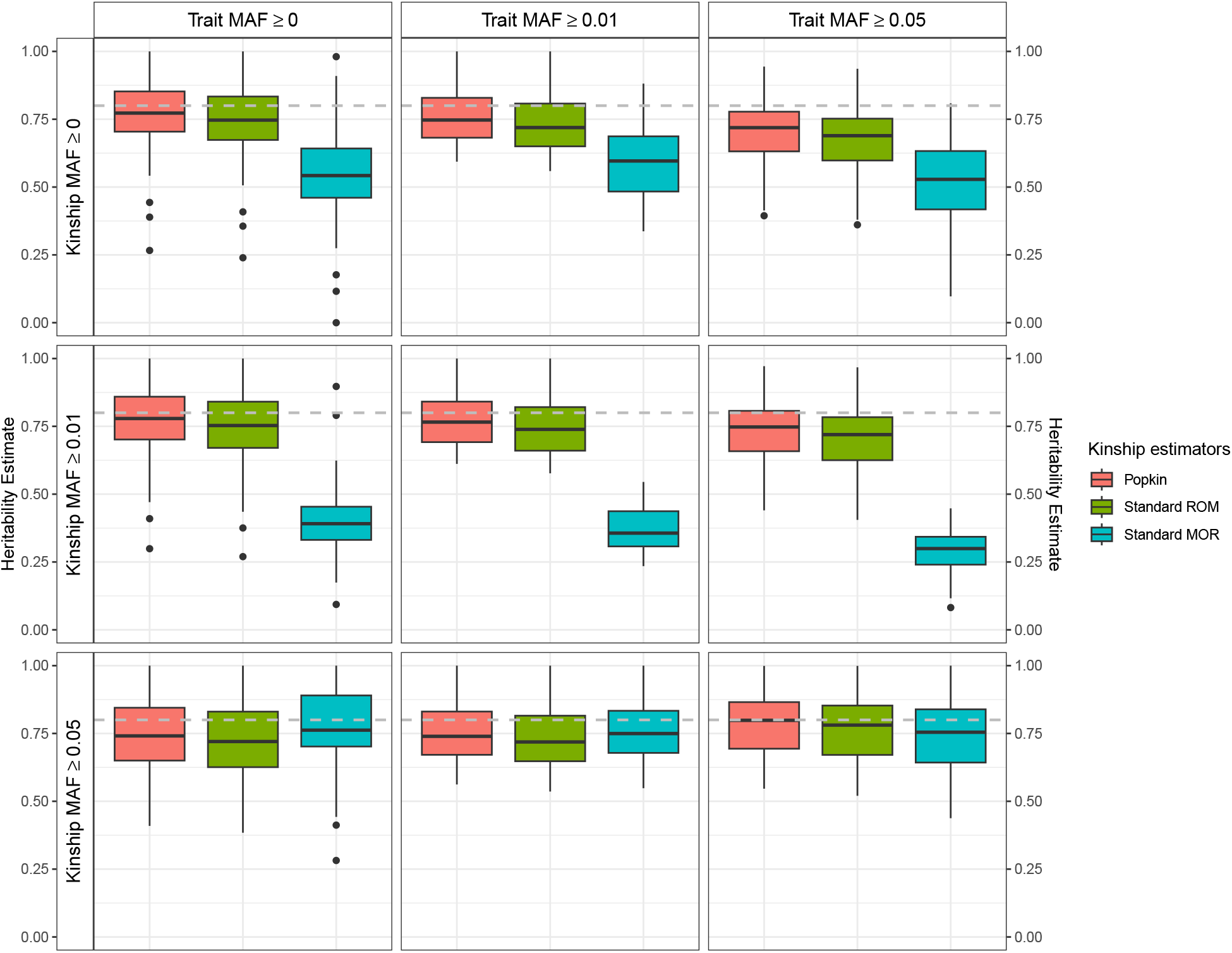
Testing effect of rare variants using 1000 Genomes genotypes and simulated traits. Here we used the RC trait model, which assigns smaller effects to causal variants and always has a correctly specified heritability. True heritability is *h*^2^ = 0.8 in all panels. Every trait simulation was replicated 50 times. For simulated traits, MAF thresholds shown are applied before causal variants are selected, thus influencing the architecture of the trait. For kinship estimates, MAF thresholds are applied to genotypes before kinship matrices are estimated from these genotypes, thus influencing only kinship estimation.

**Figure S6:**
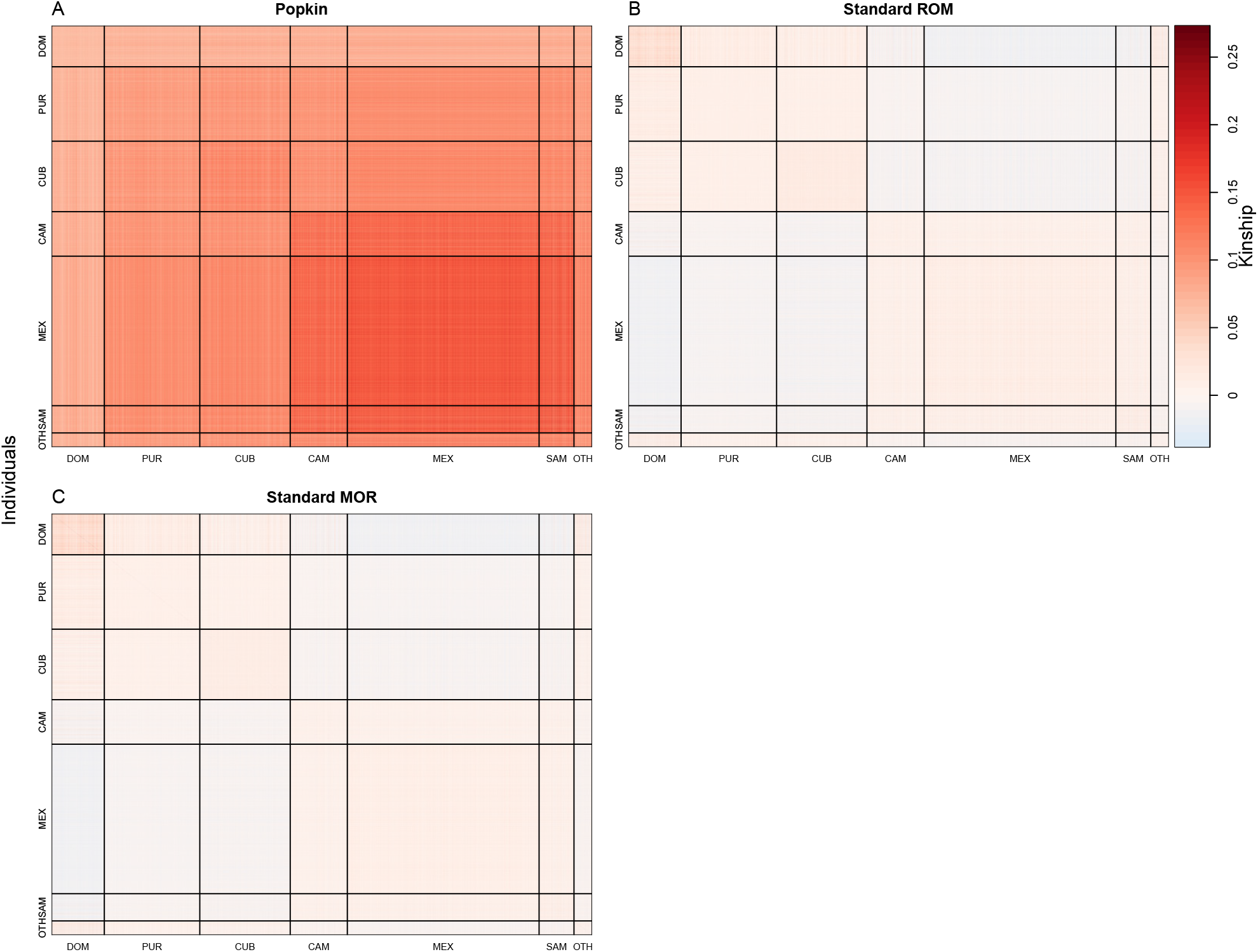
Kinship matrix estimates for the Hispanic Community Health Study / Study of Latinos dataset. DOM = Dominican, PUR = Puerto-Rican, CUB = Cuban, CAM = Central American, MEX = Mexican, SAM = South American, OTH = Other. See Fig. S1 for more details.

**Figure S7:**
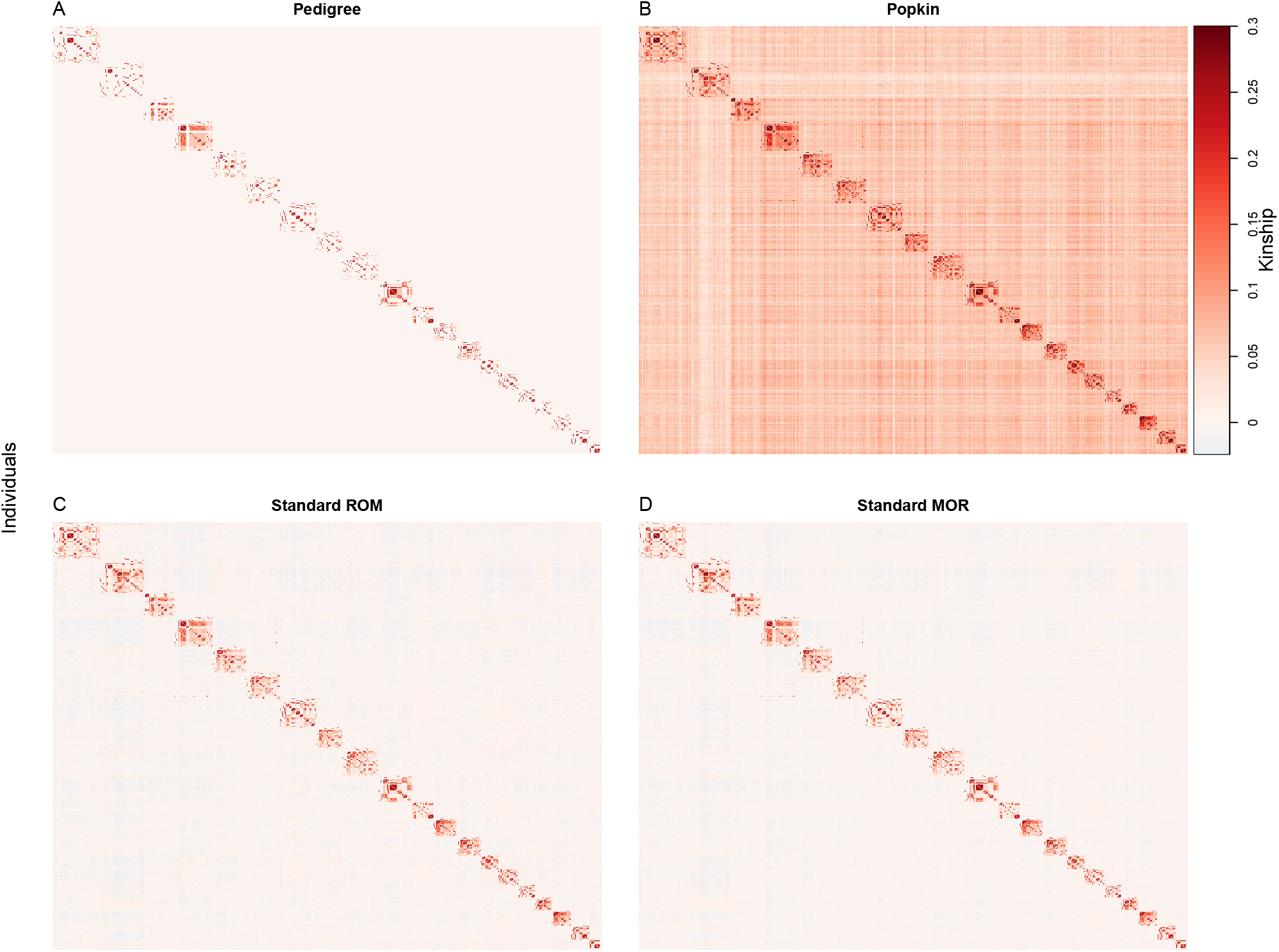
Kinship matrix estimates for the San Antonio Family Study dataset. See Fig. S1 for more details.

**Figure S8:**
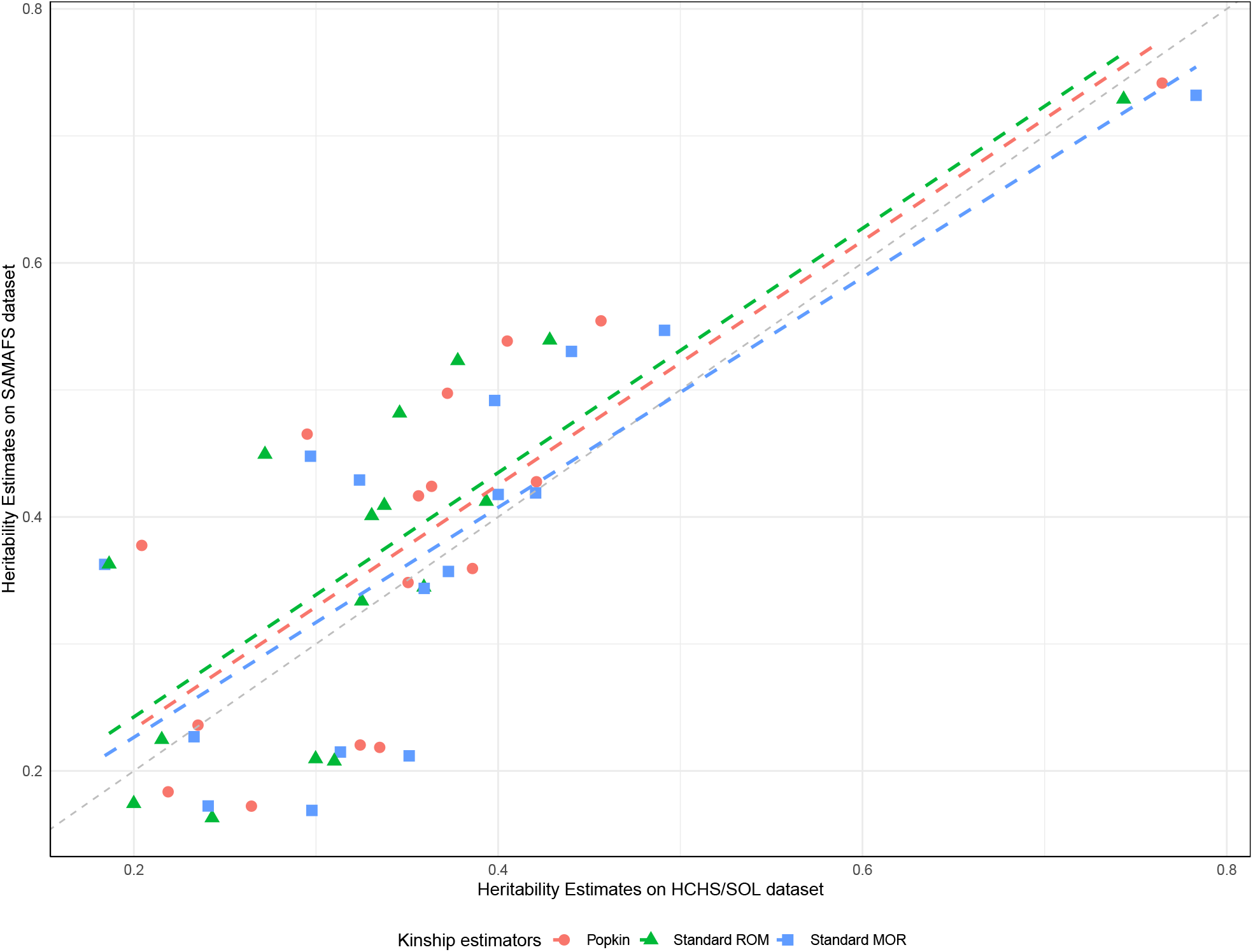
Comparison of Heritability Estimates Between HCHS/SOL and SAMAFS datasets. 16 overlapping traits are included in the analysis. Pearson correlation coefficients for each kinship estimator are: 0.799 for Popkin, 0.802 for Standard ROM, and 0.794 for Standard MOR.

**Figure S9:**
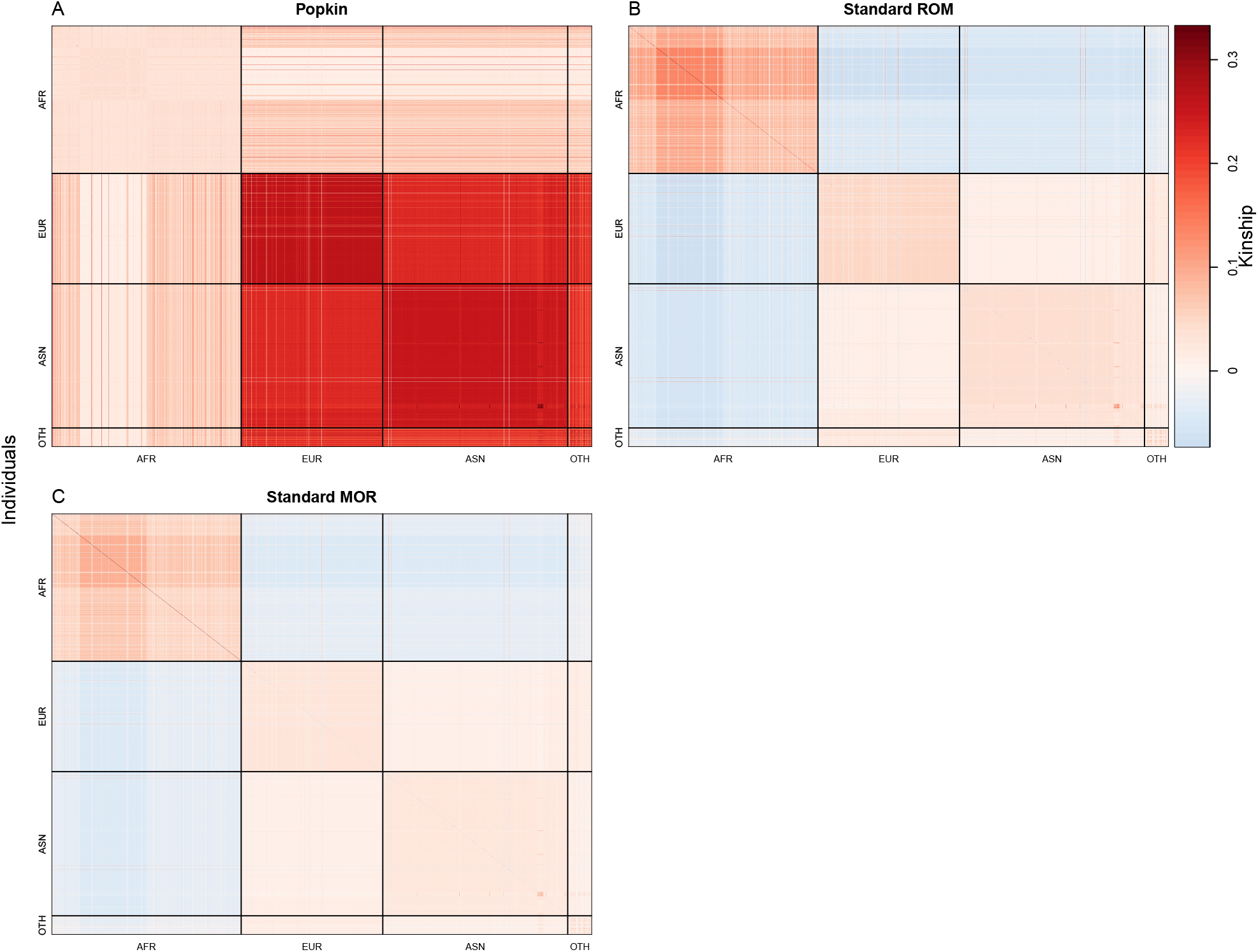
Kinship matrix estimates for the Nephrotic Syndrome multiethnic cohort dataset. AFR = African, EUR = European, ASN = Asian, OTH = Other. See Fig. S1 for more details.

